# Biochemical and structural basis of Dicer helicase function unveiled by resurrecting ancient proteins

**DOI:** 10.1101/2025.02.15.638221

**Authors:** Adedeji M. Aderounmu, Josephine Maus-Conn, Claudia D. Consalvo, Peter S. Shen, Brenda L. Bass

## Abstract

A fully functional Dicer helicase, present in the modern arthropod, uses energy generated during ATP hydrolysis to power translocation on bound dsRNA, enabling the processive dsRNA cleavage required for efficient antiviral defense. However, modern Dicer orthologs exhibit divergent helicase functions that affect their ability to contribute to antiviral defense, and moreover, mechanisms that couple ATP hydrolysis to Dicer helicase movement on dsRNA remain enigmatic. Here, we used biochemical and structural analyses of ancestrally reconstructed Dicer helicases to map evolution of dsRNA binding affinity, ATP hydrolysis and translocation. We found that loss of affinity for dsRNA occurred early in Dicer evolution, coinciding with a decline in translocation activity, despite preservation of ATP hydrolysis activity, exemplified by the ancient deuterostome Dicer. Ancestral nematode Dicer also exhibited significant decline in ATP hydrolysis and translocation, but studies of antiviral activities in the modern nematode *C. elegans* indicate Dicer retained a role in antiviral defense by recruiting a second helicase. Cryo-EM analyses of an ancient metazoan Dicer allowed capture of multiple helicase states revealing the mechanism that connects each step of ATP hydrolysis to unidirectional movement along dsRNA. Overall, our study rationalizes the diversity in modern Dicer helicases by connecting ancestral functions to observations in extant enzymes.

**Significance Statement:** Among invertebrates, the contribution of Dicer’s helicase to recognition and elimination of viral double-stranded RNA varies from phylum to phylum. At the extreme end of the spectrum, vertebrate Dicers show no helicase activity. On the other end, an arthropod ortholog uses helicase translocation to efficiently move double-stranded RNA into Dicer’s cleavage site. The biochemical and structural basis of Dicer’s helicase function, as well as the evolutionary events that contribute to a divergence in function, have remained unknown. This study shows how ancient Dicer helicase tightly binds double-stranded RNA and couples ATP hydrolysis to movement along this substrate. In addition, the data reveal how components of this intricate system declined along different clades of animal evolution.

## Introduction

Antiviral defense in vertebrates involves a family of proteins called RIG-I-like receptors (RLRs) that recognize viral dsRNA to trigger an interferon response, while invertebrates from the arthropod and nematode phyla rely on the endoribonuclease Dicer to recognize viral dsRNA and trigger antiviral RNA interference (RNAi) (1). RLRs and Dicer have closely related Superfamily 2 helicase domains, suggesting this helicase domain was involved in antiviral defense in a common ancestor (1, 2). However, helicase domains of extant Dicer enzymes have widely different activities that likely reflect an evolutionary arms race with viruses, as well as the need to acquire new functions, such as processing the microRNAs (miRNAs) required for gene regulation in modern day animals (3).

*Drosophila melanogaster* Dicer-1 (dmDcr-1) has a defunct helicase domain incapable of ATP hydrolysis. This enzyme uses its platform/PAZ domains to recognize miRNA precursors (pre-miRNAs), which are cleaved by Dicer’s RNase III domains to produce mature miRNAs (4). By contrast, dmDcr-2, encoded by a duplicated Dicer gene specific to arthropods, has a highly specialized helicase domain that couples ATP-driven dsRNA translocation to processive cleavage of long viral dsRNAs (Fig. 1A) (5–9). Humans have a single Dicer gene (hsDcr) that is specialized for miRNA production, and its helicase domain also lacks the ability to hydrolyze ATP (Fig. 1B). Phylogenetic analyses indicate this loss occurred at the onset of vertebrate evolution (10–12). Accordingly, most evidence indicates mammalian Dicers only play a role in antiviral defense in special conditions, such as in mammalian germ and stem cells, and in these cases, processing is ATP-independent (13, 14).

**Fig. 1.**
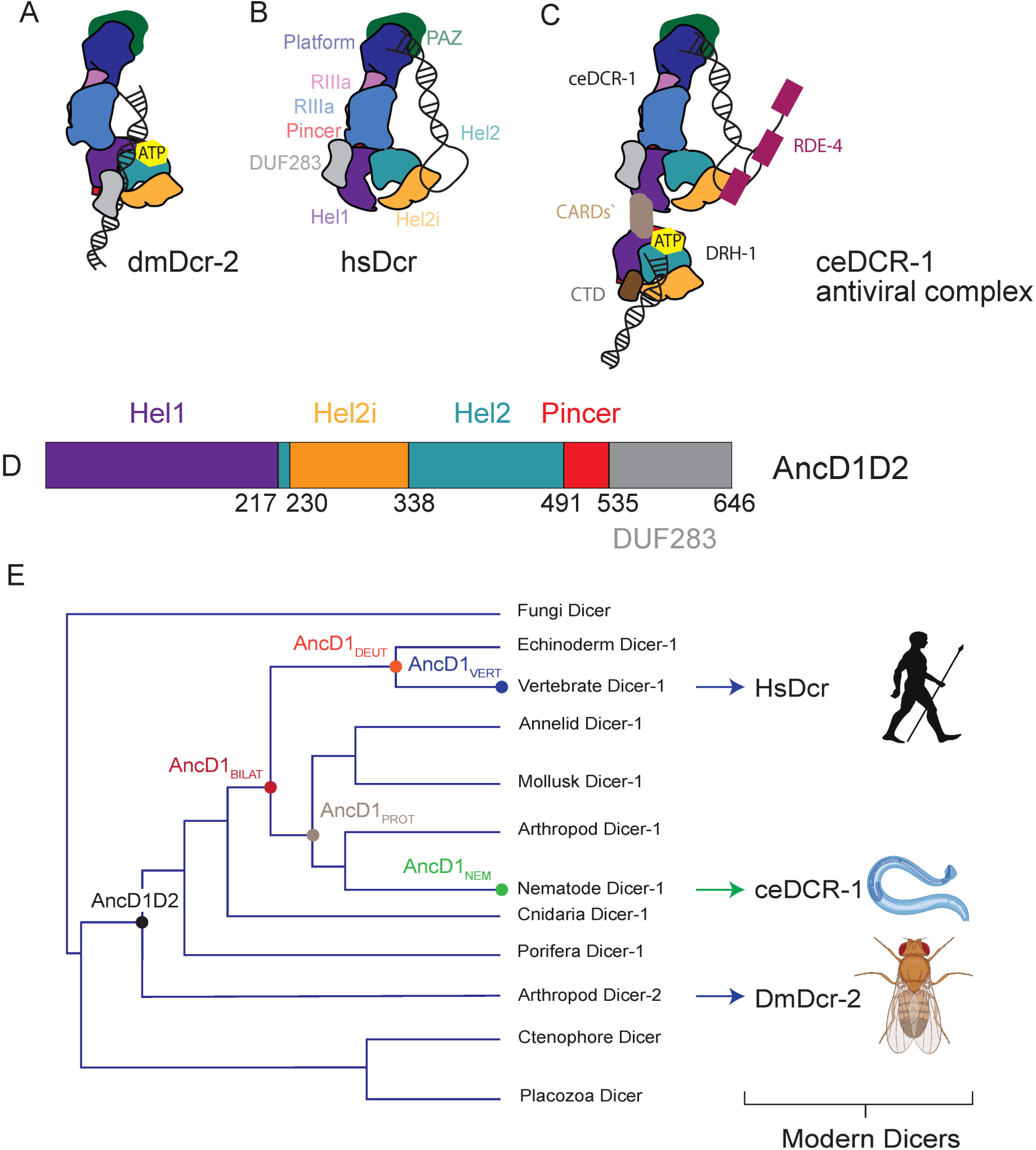
Phylogenetic analysis of diversity in Dicer helicase function. **(A-C)** Cartoon depictions of **(A)** dmDcr-2 binding long dsRNA with helicase domain, **(B)** hsDcr binding pre-miRNA with Platform/PAZ domains, and **(C)** ceDCR-1 antiviral complex binding pre-miRNA and long dsRNA using Platform/PAZ domains and DRH-1 helicase, respectively (9, 11, 20, 35). RDE-4 is shown with dsRNA binding domains (dsRBDs) in maroon. **(D)** Domain organization of AncD1D2, with colored rectangles showing conserved domain boundaries indicated by amino acid number. Domain boundaries were defined by information from NCBI Conserved Domains Database (CDD), available crystal and cryo-EM structures, and structure-based alignments from previous studies (12). **(E)** Summarized phylogenetic tree constructed by constraining maximum likelihood tree to known species relationships in Dicer-1 clade. Font colors at nodes correlate with data for ancestral protein.

The lack of helicase function in hsDcr may have arisen due to competition between helicase domains of Dicer and RLRs, as both recognize similar viral dsRNAs in the metazoan cytoplasm. In support of this idea, RNAi and RLR-stimulated interferon signaling pathways antagonize one another in mammals (15–17). Also, arthropods like *Drosophila*, where extant dmDcr-2’s helicase domain retains a role in antiviral defense, do not have RLRs. An alternative explanation is that loss of Dicer helicase function began early in animal evolution, driven by the competition for RNA substrate binding by Dicer’s helicase domain and its own Platform/PAZ domains. In this case, dominance of RLRs in viral dsRNA recognition, rather than causing loss of Dicer helicase function, would itself be an evolutionary consequence of decline in Dicer helicase function. *Caenorhabditis elegans* Dicer (ceDCR-1) functions in concert with an RLR known as DRH-1, which is crucial for dsRNA cleavage in vitro and for antiviral defense in vivo (Fig. 1C) (18–20). This raises the possibility that ceDCR-1’s helicase domain was subject to the same evolutionary pressures that led to loss of vertebrate Dicer helicase function, and that Dicer began to lose helicase function early in animal evolution, before the divergence of vertebrates and invertebrates. In this case, the hypothesis is that ceDCR-1 remedied the loss of Dicer helicase function by supplanting it with another helicase, DRH-1.

Ancestral protein reconstruction (APR) provides a powerful tool for dissecting evolution of Dicer’s helicase domain (12, 21). Previously, we used APR to track evolution of Dicer’s helicase domain in animals, mapping differences in ATP hydrolysis capabilities between arthropod and vertebrate Dicers (12). By biochemically characterizing predicted ancestral helicases, we concluded that the capacity for ATP hydrolysis was likely present in early animal Dicer at the common ancestor of arthropods and mammals but lost between common ancestors of deuterostomes and vertebrates. However, ATP hydrolysis, while an important indicator of helicase function, does not provide direct information on the motor function of Dicer’s helicase: translocation on dsRNA.

Here we extend our biochemical and phylogenetic analyses to map evolution of dsRNA translocation by Dicer’s helicase domain. This analysis, in combination with cryo-EM analyses of an ancient Dicer helicase domain, produced a more comprehensive understanding of Dicer evolution. We find that early metazoan Dicer helicase bound dsRNA with high affinity and coupled dsRNA binding with efficient ATP hydrolysis to drive robust translocation along dsRNA. However, helicase affinity for dsRNA began to decline in the Dicer-1 clade before the onset of bilaterians, declining further in deuterostome and protostome lineages. While ATP hydrolysis activity is also progressively lost along both clades, our work suggests that loss of dsRNA binding is the primary mechanism underlying the loss of translocation observed along the Dicer-1 clade. Structural analyses of ancient Dicer helicase translocating on dsRNA, likely aided by tight binding to dsRNA, provide previously unrecognized details in SF2 helicase function that are likely operative in modern RLRs.

## Results

### Ancestral protein reconstruction using alternative Dicer helicase phylogeny

The maximum likelihood (ML) phylogenetic tree generated in our prior studies fit best with a model where early metazoan Dicer (AncD1D2) underwent a gene duplication to produce Dicer-1 and Dicer-2, one of which was subsequently lost in most modern animals (12). While this model is supported by multiple independent lines of evidence in Dicer phylogenetics (21, 22), the finer details of the relationship between Dicer helicase domains defined by the ML tree did not fit the species tree of life, either due to incomplete lineage sorting or long branch attraction. This is not unexpected since the ML tree, henceforth called “gene” tree, was based on phylogenetic analyses of only two domains (helicase and DUF283, Fig 1D) of the Dicer gene, while species trees are typically built from multi-gene or whole genome datasets. Dicer gene duplications and subsequent gene loss could also explain the gene tree-species tree incongruence.

In hopes of identifying the evolutionary windows where Dicer helicase functions were lost, we repeated phylogenetic tree reconstruction using the helicase domain and DUF283 (henceforth referenced as helicase) and constrained the Dicer-1 clade of the phylogenetic tree to match evolutionary relationships from the consensus tree of life (Fig. 1E). While the species tree recapitulated some clade relationships of the gene tree, some nodes were replaced by new nodes in the species tree, providing the opportunity to perform additional analyses to gain new information (*SI Appendix*, Fig. S1A and B). Addition of the hypothetical ancestor of bilaterian Dicer1 (AncD1_BILAT_) was particularly advantageous, offering the opportunity to determine whether loss of ATP-dependent functions is unique to deuterostomes and their vertebrate descendants, or if the trend extends into invertebrate protostomes. Interestingly, the ancestral vertebrate Dicer-1 (AncD1_VERT_) primary sequence remained identical when predicted from either gene or species-constrained tree. Furthermore, as described subsequently, similar biochemical trends were observed in ancestral Dicer helicases from gene and species trees, lending confidence and robustness to the biochemical properties of ancestors predicted from phylogenies constructed with different methods.

### Ancestral animal Dicer helicase translocates on dsRNA

Previous work shows that other SF2 helicases generate enough force to displace streptavidin bound at the distal end of dsRNA (23, 24). To determine if ancient animal Dicers translocate on dsRNA, we adopted a gel-based streptavidin displacement assay that monitors the ability of translocating helicases to remove dsRNA-bound streptavidin as a measure of helicase motor function (Fig. 2A). dsRNA was ^32^P-labeled at the 5’ end of the sense (top) strand to allow visualization of radioactive dsRNA species on a native gel and conjugated with biotin at the 5’ end of the antisense (bottom) strand to allow streptavidin binding (Fig. 2B, *SI Appendix*, Fig. S2). Streptavidin was incubated with dsRNA for 20 minutes and allowed to bind conjugated biotin before addition of ATP and helicase to start the displacement reaction.

**Fig. 2.**
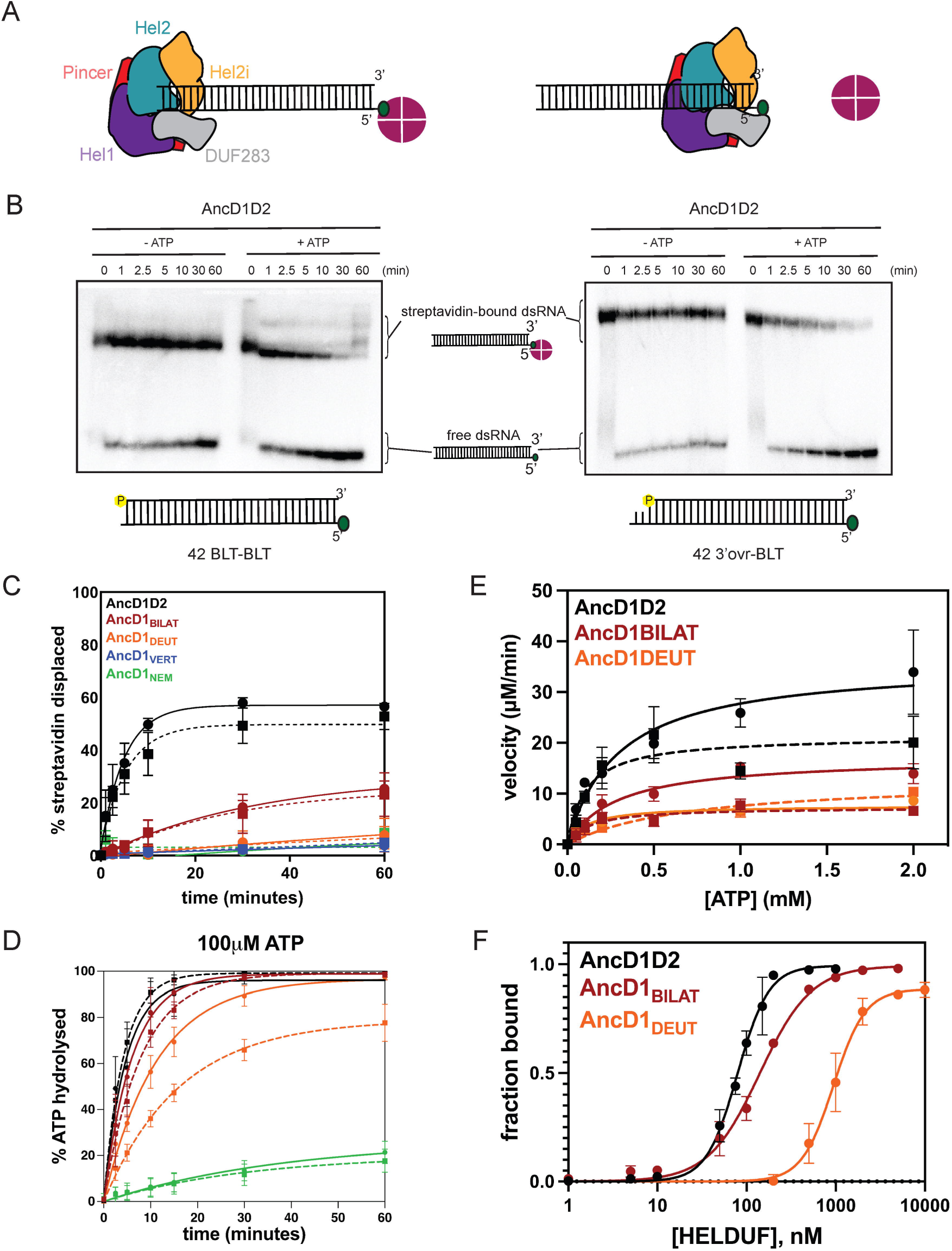
Ancient metazoan Dicer translocates along dsRNA. **(A)** Schema depicting streptavidin-displacement assay for translocation. Green oval, biotin covalently linked to RNA; Maroon circle, tetravalent streptavidin. Color-coded Dicer domains as labeled. **(B)** Representative PhosphorImages of streptavidin-displacement assays for AncD1D2 on BLT (left) and 3’ovr (right) dsRNA in the absence and presence of 5mM ATP. Yellow highlighted P, 5’ ^32^P-label. **(C)** Graph showing streptavidin displacement data for ancestral constructs from species-constrained tree for BLT (bold) and 3’ovr (dashed) 42-bp dsRNA with 5mM ATP. Data were fit to a pseudo-first order rate equation. Data points, mean ± SD (n ≥ 3). **(D)** Graph showing quantification of TLC analyses monitoring hydrolysis of ATP (100µM) by select ancestral helicases with BLT (bold) and 3’ovr (dashed) 42-bp dsRNA (400nM). Data points are mean ± SD (n ≥ 3) and were fit to a pseudo-first order equation. **(E)** Michaelis-Menten curves of ATP hydrolysis by AncD1D2, AncD1_BILAT_ and AncD1_DEUT_ with BLT (bold) and 3’ovr (dashed) dsRNA. **(F)** Binding isotherms for ancestral proteins (HEL-DUF) from gel-shift assays where fraction bound was determined from radioactivity of dsRNA_free_ and dsRNA_bound_. Data were fit to calculate dissociation constant, K_d_, using the Hill formalism, where fraction bound = 1/(1 + [K_d_^n^/[P]^n^]). Data points, mean ± SD (n≥3).

AncD1D2 displayed robust translocation on dsRNAs with blunt (BLT) and 3’overhang (3’ovr) ends (Fig. 2B and C, Table S1). With BLT dsRNA, a synthetic mimic of viral dsRNA, some ATP-independent streptavidin displacement was observed (Fig. 2B, left panel). Possibly AncD1D2 binds to ends as well as internal regions of BLT dsRNA, leading to stacking of multiple helicases, even without ATP, which in turn displaces streptavidin. Interestingly, this phenomenon was not observed with 3’ovr, the synthetic mimic of cellular pre-miRNA termini (Fig. 2B, right panel), or any other ancestral proteins. Quantification of multiple assays revealed that the bilaterian ancestor, AncD1_BILAT_, also exhibited translocation activity, albeit significantly reduced from AncD1D2, for both BLT and 3’ovr dsRNA (Fig. 2C, *SI Appendix*, Fig. S3A, Table S1). After 60 minutes, ∼30% of streptavidin had been displaced compared to 60% for AncD1D2, for both BLT and 3’ovr substrates. AncD1_DEUT_, the deuterostome descendant of bilaterian Dicer-1, showed minimal streptavidin displacement activity, with or without ATP (Fig. 2C, *SI Appendix*, Fig. S3B).

AncD1_VERT_ did not translocate on dsRNA, consistent with the absence of ATPase activity shown in prior work (Fig. 2C, *SI Appendix*, Fig. S3C)(12). A similar loss of translocation capability was observed with AncD1_NEM_, the common ancestor of the protostome Nematode Dicer-1 (Fig. 2C, *SI Appendix*, Fig. S3D). This independent loss of helicase function along two descendant lineages of AncD1_BILAT_ suggests the decline in helicase function between AncD1D2 and AncD1_BILAT_ progressed further until dsRNA translocation activity was lost in modern Dicer-1. While we did not test the ancestor of Arthropod Dicer-1, we can confidently predict loss of helicase function based on the degeneration of sequences in the Dicer-1 ATPase catalytic site in modern arthropods (4, 25).

We performed ATP hydrolysis and dsRNA binding assays to determine how these reactions underpin the observed trend in dsRNA translocation. Using a thin-layer chromatography (TLC) assay, we performed multiple-turnover ATPase assays with 100µM ATP in the presence of the ancestral protein, with and without excess dsRNA. Like the gene tree constructs (12), ancestral Dicer helicases from the species tree required dsRNA to catalyze robust ATP hydrolysis (Fig. 2D, *SI Appendix*, Fig. S4A). However, the subtle differences in hydrolysis efficiency between different Dicer ancestors did not mirror the large differences observed in dsRNA translocation between AncD1D2, AncD1_BILAT_ and AncD1_DEUT_ (Fig. 2C and D, *SI Appendix*, Fig. S4, Table S1). Attempting to resolve this incongruence, we performed Michaelis-Menten analyses on select ancestral helicases. We observed an ∼2-3-fold decrease in the efficiency of hydrolysis (k_cat_/K_M_) from AncD1D2 to AncD1_BILAT._ (Fig. 2E, *SI Appendix*, Table S1). For AncD1_DEUT_, the net k_cat_/K_M_ value was similar to AncD1_BILAT_ for BLT dsRNA but was reduced by ∼3-fold for 3’ovr (Fig. 2E, *SI Appendix*, Table S1). The preservation of ATP hydrolysis in AncD1_DEUT_ suggested the absence of translocation in AncD1_DEUT_ could not be entirely explained by a loss of hydrolysis.

Both ATP hydrolysis assays were performed with excess dsRNA compared to protein (multiple turnover conditions) to minimize contributions of dsRNA affinity to observed differences. Due to practical limitations, the translocation assay was performed with excess protein (single turnover; see Materials and Methods). To assess effects of dsRNA binding on translocation trend, we performed gel shift assays incubating AncD1D2, AncD1_BILAT_ and AncD1_DEUT_ with BLT dsRNA in the presence of ATP, using conditions similar to the translocation assay (Materials and Methods). AncD1D2 had a dissociation constant (K_D_) of 79nM compared to 140nM for AncD1_BILAT_, while AncD1_DEUT_ bound BLT dsRNA with a K_D_ of 942nM (Fig. 2F, *SI Appendix*, Fig. S5). The gel shift results suggested that the progressive decline in translocation observed from AncD1D2 to AncD1_BILAT_ to AncD1_DEUT_ is related to a loss of dsRNA affinity as the Dicer-1 helicase evolved. Thus, while ATP hydrolysis also declines along the Dicer-1 clade, our inability to detect translocation in AncD1_DEUT_ may be partially or entirely attributed to the reduction of dsRNA affinity. However, as discussed subsequently, cryo-EM analyses suggest that AncD1_DEUT_ is incapable of translocation even when bound to dsRNA. Importantly, ancestral Dicer helicases from the gene tree show a similar trend in streptavidin displacement activity. The ancient AncD1D2 and AncD1_ARTH/LOPH/DEUT_ helicases translocate along dsRNA (*SI Appendix,* Fig. S6) but this activity is progressively lost along the Dicer-1 clade, mirroring the decline in dsRNA affinity and ATP hydrolysis (12).

### *Caenorhabditis elegans* Dicer recruits an RLR, DRH-1, to perform translocation in the antiviral complex

AncD1_NEM_, the ancestor of modern ceDCR-1, displayed minimal ATP hydrolysis and translocation activity (Fig. 2C and D). ceDCR-1 functions in multiple pathways in vivo, and when targeting viral dsRNA, functions in the antiviral complex with an RLR helicase, DRH-1, and a dsRNA binding protein, RDE-4 (Fig. 1C) (18). Our recent work shows ATP hydrolysis by both helicases is required for cleavage, but that of ceDCR-1 is far less efficient than that of DRH-1 (20), consistent with the decline in hydrolysis and translocation observed in AncD1_NEM_.

Structural data in the recent study show DRH-1 localized to internal regions of dsRNA, suggesting that it fuels translocation of the antiviral complex (20). To understand how the modern *C. elegans* antiviral complex translocates, we used the streptavidin displacement assay. We first quantified translocation by the *C. elegans* antiviral complex using modern-day dmDcr-2 as a control. dmDcr-2 acts as a single protein to cleave viral dsRNA in fruit flies, and in vitro studies show it couples ATP hydrolysis to translocation (6, 26). To focus on translocation, for both organisms, we made mutations in Dicer to eliminate RNase III cleavage activity (Fig. 3A and B). Robust ATP-dependent translocation was observed for the ceDCR-1^RIII^/DRH-1/RDE-4 complex, as well as dmDcr2^RIII^, although the *C. elegans* antiviral complex was more efficient, showing similar levels of translocation at lower protein concentrations (Fig. 3A and B, *SI Appendix*, Fig. S7A and B). Consistent with prior studies of cleavage and ATP hydrolysis catalyzed by the ceDCR-1 antiviral complex and dmDcr2, translocation was more efficient with BLT than 3’ovr dsRNA (Fig. 3B) (8, 20).

**Fig. 3.**
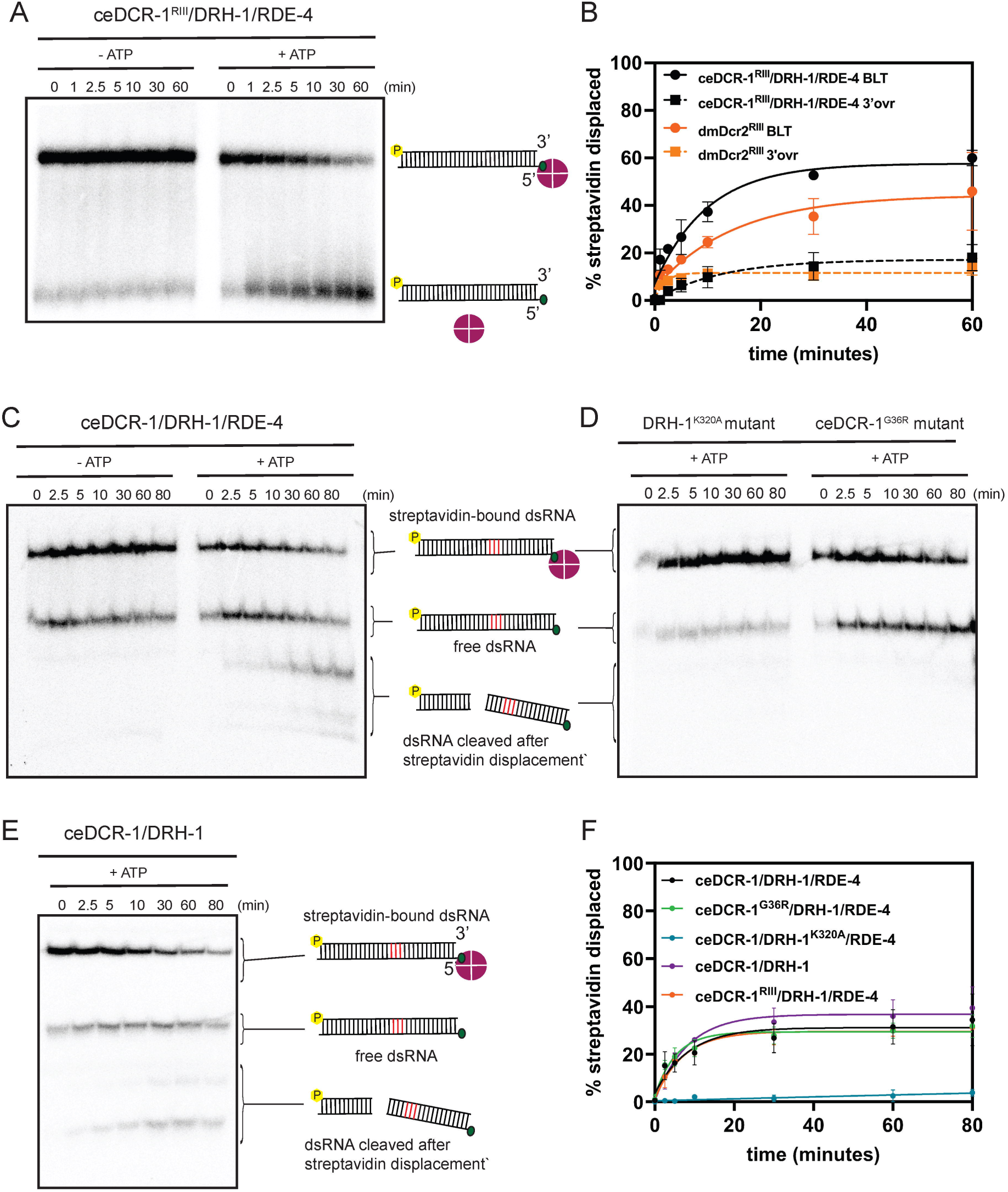
Extant invertebrate antiviral Dicers translocate along dsRNA. **(A)** Representative PhosphorImage of streptavidin-displacement assay for ceDCR-1^RIII^/DRH-1/RDE-4 with BLT 42-bp dsRNA, with and without 5mM ATP. **(B)** Graph quantifying streptavidin displacement data for 50nM ceDCR-1^RIII^/DRH-1/RDE-4 and 200nM dmDcr2^RIII^, for BLT and 3’ovr dsRNA with 5mM ATP. Data were fit to a pseudo-first order rate equation. Data points, mean ± SD (n = 3). **(C)** Representative PhosphorImage of streptavidin-displacement assay for ceDCR-1/DRH-1/RDE-4 with BLT 42-bp dsRNA with deoxynucleotide patch, with and without 5mM ATP. Cartoons indicate deoxynucleotides (red) and gel migration of different species. **(D)** Representative PhosphorImage of streptavidin-displacement assay for ceDCR-1/DRH-1^K320A^/RDE-4 and ceDCR-1^G36R/^DRH-1/RDE-4 with BLT 42-bp dsRNA with deoxynucleotide patch with 5mM ATP. Cartoons indicate deoxynucleotides (red) and gel migration of species. **(E)** Representative PhosphorImage of streptavidin-displacement assay for ceDCR-1/DRH-1 with BLT 42-bp dsRNA containing deoxynucleotides with 5mM ATP. Cartoons indicate deoxynucleotides (red) and gel migration of species. **(F)** Graph quantifying streptavidin displacement data for ceDCR-1/DRH-1/RDE-4 and indicated mutants, using BLT 42-bp BLT dsRNA with deoxynucleotides and 5mM ATP. Data were fit to a pseudo-first order rate equation. Data points, mean ± SD (n = 3).

To delineate contributions of each protein of the antiviral complex to translocation, we designed a version of the assay that did not require RNase III mutation, using 42-bp dsRNAs with deoxynucleotides at predicted ceDCR-1 cleavage sites (*SI Appendix*, Fig. S2 and S8). Wildtype *C. elegans* antiviral complex showed ATP-dependent streptavidin displacement on this substrate, with minimal dsRNA cleavage (Fig. 3C). A point mutation in the Walker A motif of DRH-1’s helicase domain (DRH-1^K320A^) abolished all translocation activity, while mutating the Walker A motif in ceDCR-1 (ceDCR-1^G36R^), or omitting RDE-4 from the complex, did not affect translocation (Fig. 3D-F). Thus, DRH-1, not ceDCR-1, is responsible for translocation by the antiviral complex. While ceDCR-1 helicase mutant retained streptavidin displacement activity, it did not cleave dsRNA (compare Fig. 3C and D), suggesting that ceDCR-1’s helicase retains an ATP-dependent activity important for cleavage (27).

### AncD1D2 undergoes ATP-dependent conformational changes that are coupled to translocation on dsRNA

To understand how ancient Dicer helicase used ATP hydrolysis to power movement on dsRNA, we determined cryo-EM structures of AncD1D2 bound to a 27-bp BLT dsRNA in the absence of nucleotide, with ADP-aluminum fluoride (ADP-AlFx), and with ATP, at resolutions between 3.0-3.5 Å (Fig. 4, *SI Appendix*, Fig. S9-13, Table S2). Image processing recovered five structural snapshots: state A^0ground^ (no nucleotide, ground state), state B^end‡^ (endbound transition state), state C^post,closed^ (post hydrolysis closed state), state D^post,ground^ (post hydrolysis ground state) and state E^internal‡^ (internal transition state) (Fig. 4). As described subsequently, each state represented a unique conformation of the helicase as it cycles through a complete ATP hydrolysis reaction and couples this hydrolysis to motor function. Consistent with prior models (28), we observed the tandem RecA domains, Hel1 and Hel2, wrapped around dsRNA in a C-shaped enclosure in concert with the C-terminal dsRBM fold, DUF283, which is connected to Hel2 by a pincer domain that regulates helicase function in related helicases (Fig. 4A-E, far left column).

**Fig. 4.**
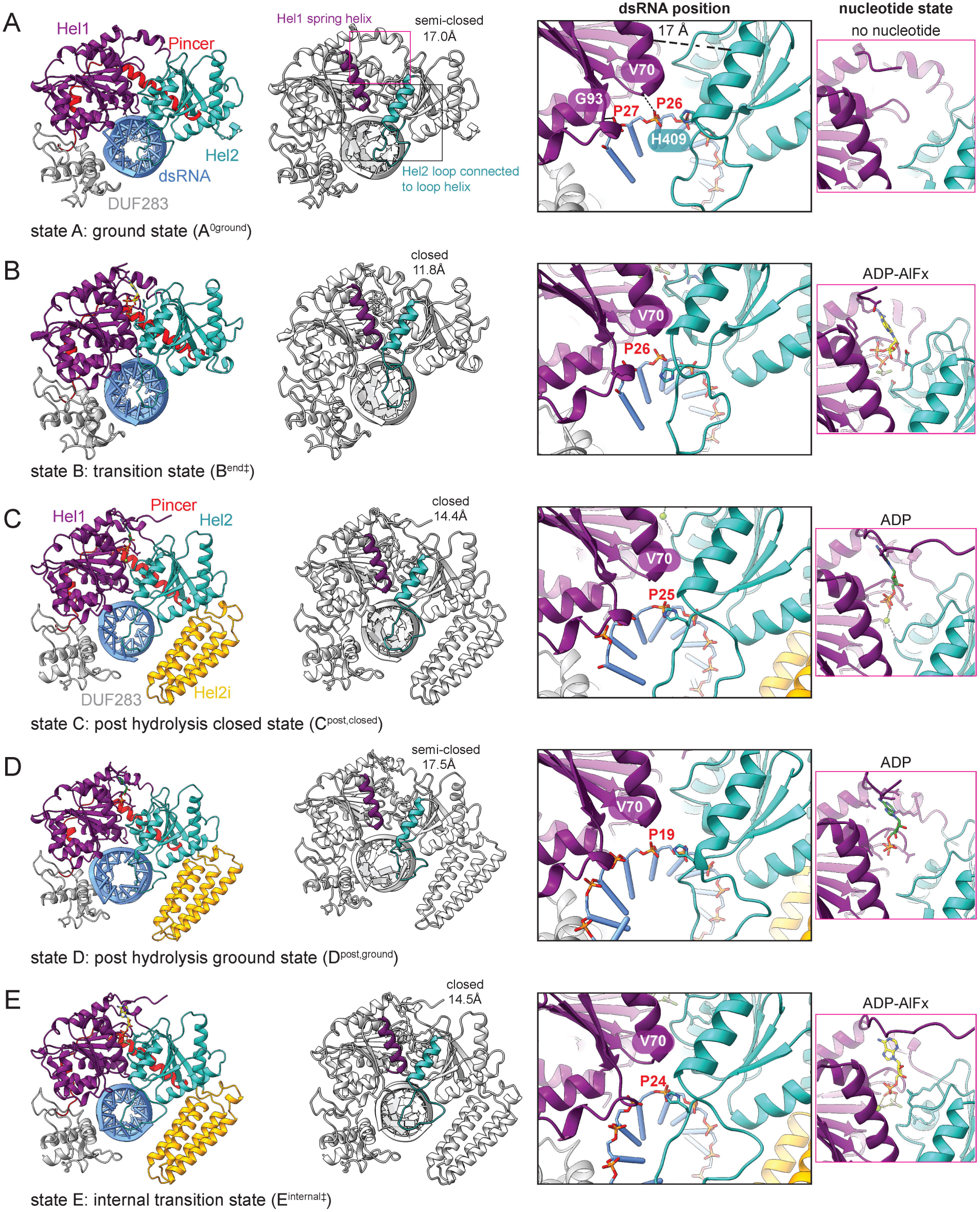
Cryo-EM structures of AncD1D2 in complex with BLT 27-dsRNA in different stages of ATPase cycle. Colored atomic models of AncD1D2 in complex with BLT 27-dsRNA in **(A)** state A^0ground^, ground state without nucleotide, **(B)** state B^end‡^, transition state following ADP-AlFx binding to ground state, **(C)** state C^post,closed^, post hydrolysis closed state following state B^end‡^, **(D)** state D^post,ground^, post hydrolysis semi-closed state in internal dsRNA segment following state C^post,closed^, **(E)** state E^internal‡^, transition state after initial ATPase cycle at dsRNA terminus. Hel2i is not visible in states A^0ground^ and B^end‡^ likely due to subdomain flexibility. Colors: Hel1 (purple), Hel2 (sea green), Hel2i (yellow), Pincer (red), DUF283 (dark gray), dsRNA (cornflower blue). Distance across ATPase cleft calculated by measuring distance between central point of Hel1 spring helix and Hel2 loop helix. dsRNA position column shows zoom-in of contacts between spring and loop helices and dsRNA 3’ tracking strand. Polar contact between V70 from spring helix, and labeled PO_4_ group in state A, was used to benchmark helicase position on dsRNA for all states. Nucleotide state column shows ATPase pocket and occupying nucleotide.

Hel2i, an SF2 helicase-specific insertion between Hel1 and Hel2, was not visible in states A^0ground^ and B^end‡^, likely due to flexibility, but was observed in states C-E. As AncD1D2 progresses through the ATPase cycle, the alpha helix in Hel1 that contains motif Ia, termed the spring helix in a related SF2 helicase (29), and the Hel2 helix containing motif IVa, termed the loop helix for its connection to the RLR helicase-specific Hel2 loop (30), moved relative to each other to switch between semi-closed and closed conformations (Fig. 4A-E, second column) (31, 32).

In state A^0ground^, AncD1D2 was bound near the dsRNA terminus with backbone amines of G93 of motif Ib and V70 of motif Ia contacting phosphate oxygens of the 3’ terminal P27 and P26 respectively (Fig. 4A, dsRNA position column). In B^end‡^, C^post,closed^ and E^internal‡^, V70 was shifted, but still interacting near the dsRNA terminus, while D^post,ground^, was positioned at an internal dsRNA segment, with V70 contacting P19 (Fig. 4A-E, dsRNA position column). Combining information from the position of AncD1D2 on dsRNA, the relative positions of spring and loop helices, and the orientation of nucleotide in the ATPase pocket (nucleotide state column), allowed us to correlate each structural snapshot with stages of the ATPase cycle (Fig. 4A-E), as described in the next section.

### ATP-stimulated closure of the helicase is the first step in AncD1D2 translocation

In state A^0ground^, Hel1 uses motifs Ia and Ib while Hel2 uses motifs IVa, IVb and V, to contact the end of the 27-bp dsRNA with an ∼8-bp footprint (Fig. 5A, *SI Appendix*, Fig. S9A). To interpret helicase movement along dsRNA as AncD1D2 cycles through ATP binding, hydrolysis and product release steps, we assigned initial Hel1-bound 3’ strand phosphates as P1 and P2. In A^0ground^, motif Ib (92-V**G**DMD-96) binds the first 3’ phosphate (P1, equivalent to P27 in Fig. 4) and motif Ia (68-NT**V**-70) binds the adjacent phosphate (P2) while motif IVa (406-IVG**H**-409) binds P3 (Fig. 5A). Motif V contacts the next phosphate and ribose sugar (*SI Appendix*, Fig S9A). This contiguous interaction between the helicase domain and the first four phosphates on the 3’ strand is the ground state configuration prior to addition of ATP. In this conformation, the ATPase cleft between Hel1 and Hel2 is in the semi-closed state, as determined by the distance separating the Hel1 spring helix and Hel2 loop helix (Fig. 5A).

**Fig. 5.**
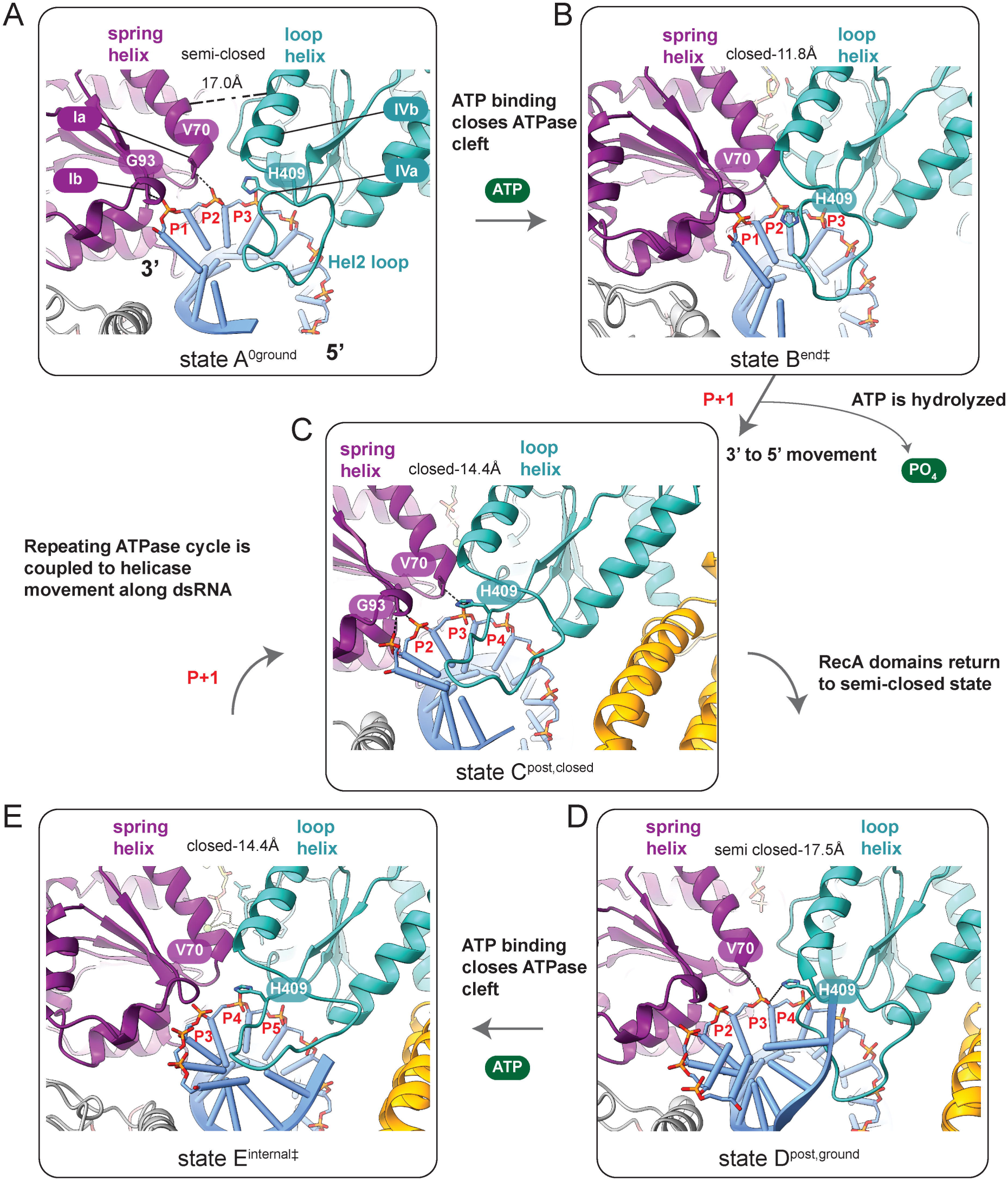
Chronology of AncD1D2 ATPase cycle coupled to translocation on dsRNA tracking strand. **(A-E)** Zoom-in of RecA-dsRNA interface, showing select Hel1 and Hel2 motifs in contact with dsRNA 3’ tracking strand for states A-E. Hel1, purple; Hel2, sea green; dsRNA, cornflower blue. Phosphates (red and yellow) connect RNA bases (tubes). Key residues are shown in spring and loop helices. Motifs are labeled and highlighted with lines depicting position in helicase. Phosphate position is benchmarked by assigning PO_4_ group contacting V70A in state A^0ground^ as P2, and correlating helicase movement on dsRNA to stages of ATPase cycle, as determined by nucleotide state in ATPase pocket and state of ATPase cleft. Interactions between protein and RNA, dashed lines. Distance (in Angstroms) between center of spring helix and center of loop helix was used to depict semi-closed or closed state. Arrows represent chronological transition from state to state.

We added ATP, NaF and Al(NO_3_)_3_ to generate ADP-AlFx and mimic the transition state of a helicase-bound ATP. This led to state B^end‡^ and showed closure of the ATPase cleft around the ATP analog, switching the helicase from the ground state to the transition state (compare Fig. 5A with 5B). The spring and loop helices moved closer together to bridge ATP, causing Hel1 to move 3’ to 5’ to form the high energy transition state (Fig. 5A and B, *SI Appendix*, Fig. S9A and B). V70 and H409 moved into proximity in the transition state where they both contact P2 (Movie S1).

### ATP hydrolysis is coupled to helicase movement along 3’ tracking strand

For Dicer’s helicase to translocate, it needs to progress beyond the transition state by coupling the ATP hydrolysis step, and subsequent product release, to motor function that results in net 3’ to 5’ movement. To visualize active translocation, we incubated AncD1D2 with 27-bp BLT dsRNA and ATP followed by incubation at 37°C for 5-10 minutes. We solved multiple cryo-EM structures that showed AncD1D2 in internal dsRNA segments with ADP in the ATPase pocket indicating a post hydrolytic stage. Focused 3D classification of this dataset yielded a unique class density depicting a helicase conformation close to the dsRNA terminus, C^post,closed^, where Hel1 and Hel2 remained in the closed conformation and contained ADP in the ATPase cleft. V70 and H409 contact P3 instead of P2 indicating movement of the helicase by one nucleotide in the 3’ to 5’ direction compared to B^end‡^ (Fig. 5C, *SI Appendix*, Fig. S9C and Fig. S12A). The presence of ADP, not ATP, indicates that C^post,closed^ represents the helicase conformation following removal of the scissile γ-PO_4_ from ATP. C^post,closed^ is a structural snapshot not previously observed, for the first time showing that the helicase moves in the 3’ to 5’ direction while the ATPase cleft is still closed (Movie S2). This observation offers insight relevant to other dsRNA-stimulated SF2 helicases.

Interestingly, multiple attempts to capture AncD1_DEUT_ in the active translocation state yielded 2D classes with density for the helicase exclusively at the dsRNA terminus. Orientation bias of the frozen sample precluded construction of high-resolution 3D density maps, but 2D classes support the biochemical assays and show a lack of translocation despite reaction conditions promoting dsRNA terminus binding (*SI Appendix*, Fig. S14). All visible particles show helicase density at the end of dsRNA, with no particles internally (*SI Appendix*, Fig. S14). Possibly AncD1_DEUT_ is missing an essential helicase component that enables coupling of ATP hydrolysis to translocation or is incapable of sustaining binding to the internal dsRNA stem and dissociates rapidly upon transition from terminal to internal dsRNA.

### AncD1D2’s transition from closed to semi-closed completes translocation along one base-pair

The remaining 3D classes from the post hydrolysis dataset show AncD1D2 in internal dsRNA regions (*SI Appendix*, Fig. S9D and Fig. S12A). The consensus model, with all subdomains visible, is D^post,ground^ (Fig. 5D, *SI Appendix*, Fig. S12A). In D^post,ground^, the spring and loop helices are in the semi-closed conformation of the ground state, despite the presence of ADP in the ATPase pocket (Fig. 5D, *SI Appendix*, Fig. S9D). This observation suggests progression from C to D is the final stage of the ATPase cycle, relaxation to the ground state. The return of Hel1 and Hel2 to the ground state changes contacts between spring and loop helices, and the dsRNA tracking strand (Fig. 5C and D). In C^post,closed^, H409 sidechain (IVa) contacts the same P3 backbone phosphate as V70, but in D^post,ground^, H409 has drifted away from P3 in the 3’ to 5’ direction (Fig. 5D, Movie S3). Hel1, now disconnected from Hel2 may pull on P3 as it relaxes in the 5’ to 3’ direction, while Hel2 could passively drift back to its ground state in the opposite direction.

To further visualize impact of ATPase pocket closure on Hel1 and Hel2, we solved a 3.1Å cryo-EM structure of AncD1D2 bound to ADP-AlFx in E^internal‡^, a state that depicts the start of the second ATPase cycle (Fig. 5E, *SI Appendix*, Fig. S9E). In E^internal‡^, the helicase returns to the closed conformation of the transition state as a new ATP mimic is bound to initiate a second round of hydrolysis and continue translocation along dsRNA. V70 and H409 are again in position to bind the same phosphate, this time the adjacent P4 phosphate.

Comparison of D^post,ground^ subclasses showed the Hel2 loop adopting different positions, while the remainder of the helicase remained roughly identical (*SI Appendix*, Fig. S12A and S15A-B). These Hel2 loop movements caused changes in contacts made by H409 in different state D subclasses (Movie S4, *SI Appendix*, Fig. S15A-B). The Hel2 loop also makes contacts with the Hel2i bundle via salt bridges in proximity to conserved residues segments (*SI Appendix*, Fig. S15C-D) and widens the dsRNA major groove as it transitions from the dsRNA terminus to internal segments (*SI Appendix*, Fig. S16A-B). Bending of the dsRNA because of the major groove expansion enables extensive contact between the internal dsRNA stem and DUF283 that may contribute to AncD1D2 translocation on dsRNA or binding affinity to internal dsRNA stem (*SI Appendix*, Fig. S9C and S16C-D).

### Closure of the ATPase cleft creates contacts between Hel1 and Hel2 that enable translocation

The higher quality of the E^internal‡^ electron density map allowed closer analysis of AncD1D2 RecA domain movement during ATP hydrolysis. Comparing B^end‡^ and C^post,closed^ suggested that the two RecA domains move as one rigid body during or immediately after ATP γ-PO_4_ bond cleavage (Fig. 5B to Fig. 5C), and we were interested in interactions that allowed this closed conformation to be maintained during translocation. Examination of the AncD1D2 ATPase pocket in E^internal‡^ confirmed that the AlFx mimic of the γ-PO_4_ is positioned proximal to the arginine finger of motif VI, while catalytic E143 from the D**E**CH-box is poised to coordinate the nucleophilic water for attack on the ß-γ PO_4_ bond (Fig. 6A, left). Closure of the ATPase cleft created a network of polar interactions between Hel1 and Hel2 (Fig. 6A, right panel). Q75 from Hel1, adjacent to Q76, contacts D453 from motif Va in Hel2, which is proximal to the ADP ribose sugar. E450 (Va), forms a network of contacts between H146 (II), S177, N180 and R470. S177 and N180 were adjacent to motif III while R470 was adjacent to motif VI. In C^post,closed^, hydrolysis of the γ-PO_4_ disrupts connection between the nucleotide and Hel2; D453 (Va) and R480 (VI) sidechains adopted conformations unable to bind ADP (Fig. 6B, left). Also, Q76 was no longer in position to coordinate the magnesium ion. However, the network of interactions between Hel1 and Hel2 was largely maintained, causing the helicase to remain in the closed state (Fig. 6B, right). These electrostatic interactions explain why the helicase remains closed in C^post,closed^ despite the presence of ADP which is unable to bridge the RecA domains.

**Fig. 6.**
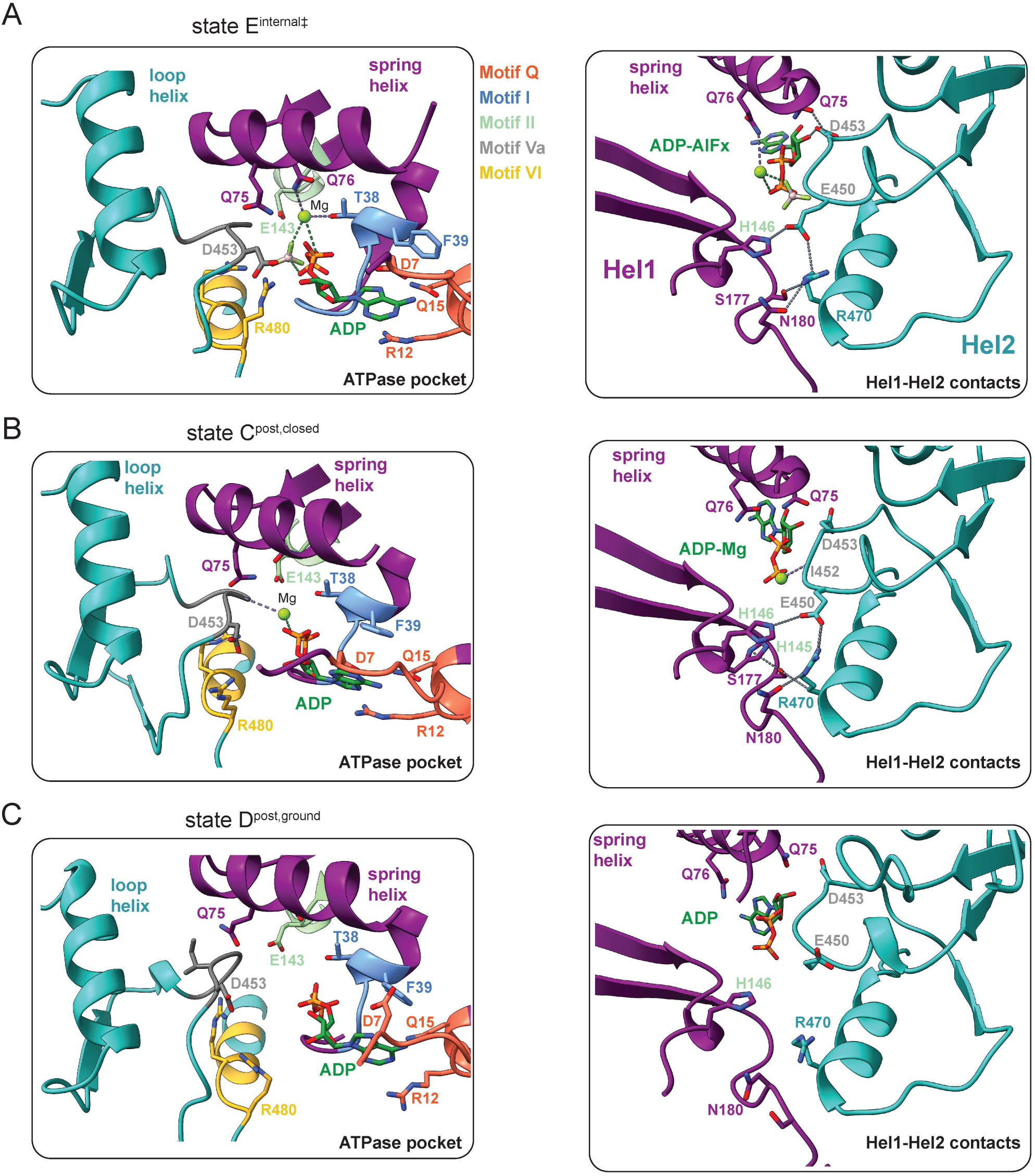
Contact between Hel1 and Hel2 is mediated by closure of ATPase pocket. Zoom-in of ATPase pocket (left) and Hel1-Hel2 interface (right) for **(A)** E^internal‡^, **(B)** C^post,closed^ and **(C)** D^post,ground^. Hel1, purple; Hel2, sea green. Select sidechains colored according to motif. Left panel: Sidechains important for ATPase function. ADP, magnesium ion and aluminum-fluoride are shown. Right panel: Side chain interactions between Hel1 and Hel2 interface residues represented by dashed lines. Non-ATPase pocket sidechains are colored according to subdomains Hel1 and Hel2.

In the transition between C^post,closed^ and D^post,ground^, the network breaks down as Hel2 drifts away from Hel1. We observed D453 (Va) and R480 (VI) sidechains further away from ADP (Fig. 6C, left). The sidechains of E453 (Va) and R470 adopted positions unable to support the connection between Hel1 and Hel2 (Fig. 6C, right). This order of events suggests that formation of these contacts during closure of the RecA domains enables the helicase to move as a single body during the hydrolysis step of the ATPase cycle. Subsequent disruption of these contacts coincides with Hel1 and Hel2 individually returning to the semi-closed, ground state, adopting a conformation primed for the next round of ATP binding and hydrolysis. This network of interactions is conserved in RIG-I, supporting the notion that it mediates contact between RecA domains in the RLR family of SF2 helicases (*SI Appendix*, Fig. S17).

## Discussion

Using Ancestral Protein Reconstruction (APR) we predicted amino acid sequences of key ancestors in the evolution of the animal Dicer helicase domain, purified the ancestral proteins, and assayed dsRNA binding affinity, ATP hydrolysis and translocation along dsRNA. The common ancestor, AncD1D2, exhibited robust activity in all assays, but helicases of the Dicer-1 clade showed significant decline in all activities during evolution. An advance to our prior studies was the reconstruction of the hypothetical bilaterian ancestor (Fig. 1E), which showed a decrease of all activities, thus inferring loss of activity in both deuterostome and protostome descendants. While the AncD1_VERT_ descendent of deuterostomes showed loss of all helicase activities, the nematode descendent of protostomes, AncD1_NEM_, showed minimal ATP hydrolysis and undetectable translocation. Our studies show that a modern-day solution to loss of translocation in AncD1_NEM_ is offered by the *C. elegans* antiviral complex, which supplanted this activity by co-opting the translocation-competent RLR, DRH-1. While we observed a decrease in all helicase activities as Dicer-1 evolved, a clear driving force was loss of dsRNA affinity. Indeed, tight binding of the common ancestor, AncD1D2, may have allowed us to trap structural conformations that are likely at play in modern helicase domains, but recalcitrant to isolation.

### Biochemical properties of ancient Dicer-1 helicases explain diversity in modern helicase function

Insights from biochemical analyses of ancient Dicer helicases serve to rationalize the sequence and functional diversity observed in extant Dicers. Subfunctionalization of Dicer-1’s helicase domain in non-arthropod bilaterians was likely a stochastic process that began with progressive loss of dsRNA binding and translocation, while ATP hydrolysis remained relatively intact (Fig. 2). The delay in complete loss of ATP hydrolysis in deuterostomes, which in our studies was not observed until AncD1_VERT_, (12), explains why hsDcr retained sequence conservation in motifs responsible for ATP binding and hydrolysis despite loss of hydrolysis. The decline in dsRNA binding and translocation may have been sufficient to render the Dicer helicase defunct as a competitor for viral dsRNAs with ancient RLRs. This in turn would remove the selection pressure for further helicase subfunctionalization and allow retention of vestigial ATP hydrolysis up to AncD1_DEUT_, before it is entirely lost in AncD1_VERT_.

In the protostome clade, a minimal amount of ATP hydrolysis is retained by the AncD1_NEM_ helicase (Fig. 2D). Possibly this ATP hydrolysis is vestigial, but it clearly serves a function in modern ceDCR-1 where it is required for proper dsRNA cleavage (27), but not translocation (Fig. 3). Conceivably, ATP hydrolysis by ceDCR-1 is required to connect DRH-1 mediated translocation to dsRNA cleavage by ceDCR-1. In the Dicer-1 arthropod lineage, the ATPase pocket is degenerate, making it unlikely that hydrolysis or translocation is retained in the presence of the functional duplicate arthropod Dicer-2. Thus, the stochastic nature of Dicer-1’s evolution is reflected in diverse modern adaptations to loss of helicase function.

### Mechanistic basis of AncD1D2 translocation along dsRNA

Variation in the helicase motor function of translocation is governed by the capacity of ancestral proteins to bind dsRNA, hydrolyze ATP, and couple these activities to movement. We observed that AncD1D2 translocates efficiently on dsRNA, but this activity starts to decline in AncD1_BILAT_, concomitant with a decrease in dsRNA affinity and ATPase turnover (Fig. 2C-F). While AncD1_DEUT_ exhibited reduced levels of ATP hydrolysis, translocation in the streptavidin displacement assay was undetectable, likely due to its very low dsRNA binding affinity (Fig. 2C-F). However, our cryo-EM studies were performed under saturating dsRNA conditions and revealed AncD1_DEUT_ at the dsRNA terminus, but not in internal regions (*SI Appendix*, Fig. S14). Thus, further decline in translocation capability appears to have been caused by the inability of AncD1_DEUT_ to move into dsRNA, despite preservation of its ability to hydrolyze ATP (Fig. 2D and E).

Our structural analyses of AncD1D2 helicase in multiple states (Fig. 4-6) provide information on helicase function that can be extended to modern Dicers and RLRs (Fig. 7A). Structural studies on extant dmDcr-2 showed that dsRNA binding to its helicase domain, in the absence of ATP, causes a conformational change from the open to the semi-closed state (9). The ground state of AncD1D2, A^0ground^, captured in the absence of nucleotide, exists in a similar semi-closed state bound to the dsRNA terminus (Fig. 4A). ATP binding caused Hel1 and Hel2 RecA domains to close further to form a high-energy transition state observed in both endbound (B^end‡^) and internally bound (E^internal‡^) AncD1D2 (Fig. 5A and E and 7B, Movie S1). Closed transition states have been observed in RLR helicases containing transition state mimics (31, 33) and in AAA+ helicases (34), but the mechanism that governs the coupling of ATP binding and hydrolysis in Dicer to successive transitions between semi-closed and closed states, and the resulting motor function of unidirectional translocation, has remained enigmatic. By isolating C^post,closed^, the post hydrolytic closed state, and D^post,ground^, the post hydrolytic semi-closed state, we find that movement of the helicase on dsRNA occurs in the closed state, concurrent with the ATP hydrolysis step. Conformational changes that occur in Hel2 as the γ-PO_4_ is cleaved and dislodged from motif VI drive movement of Hel2 in the 5’ direction while interactions between Hel2 and Hel1 ensure that Hel1 is pulled in the same direction (Fig. 6A). Our cryo-EM snapshots depict 3’ to 5’ movement as the transition from B^end‡^ to C^post,closed^, which is then followed by relaxation from C^post,closed^ to D^post.ground^ (Fig. 5B-D, Movie S2, S3). Structural analysis of an ancient Dicer helicase thus fills in the gaps in the second half of the ATPase cycle, revealing the movement of closed helicase on dsRNA followed by the relaxation of the closed post hydrolysis state to the semi-closed ground state.

**Fig. 7.**
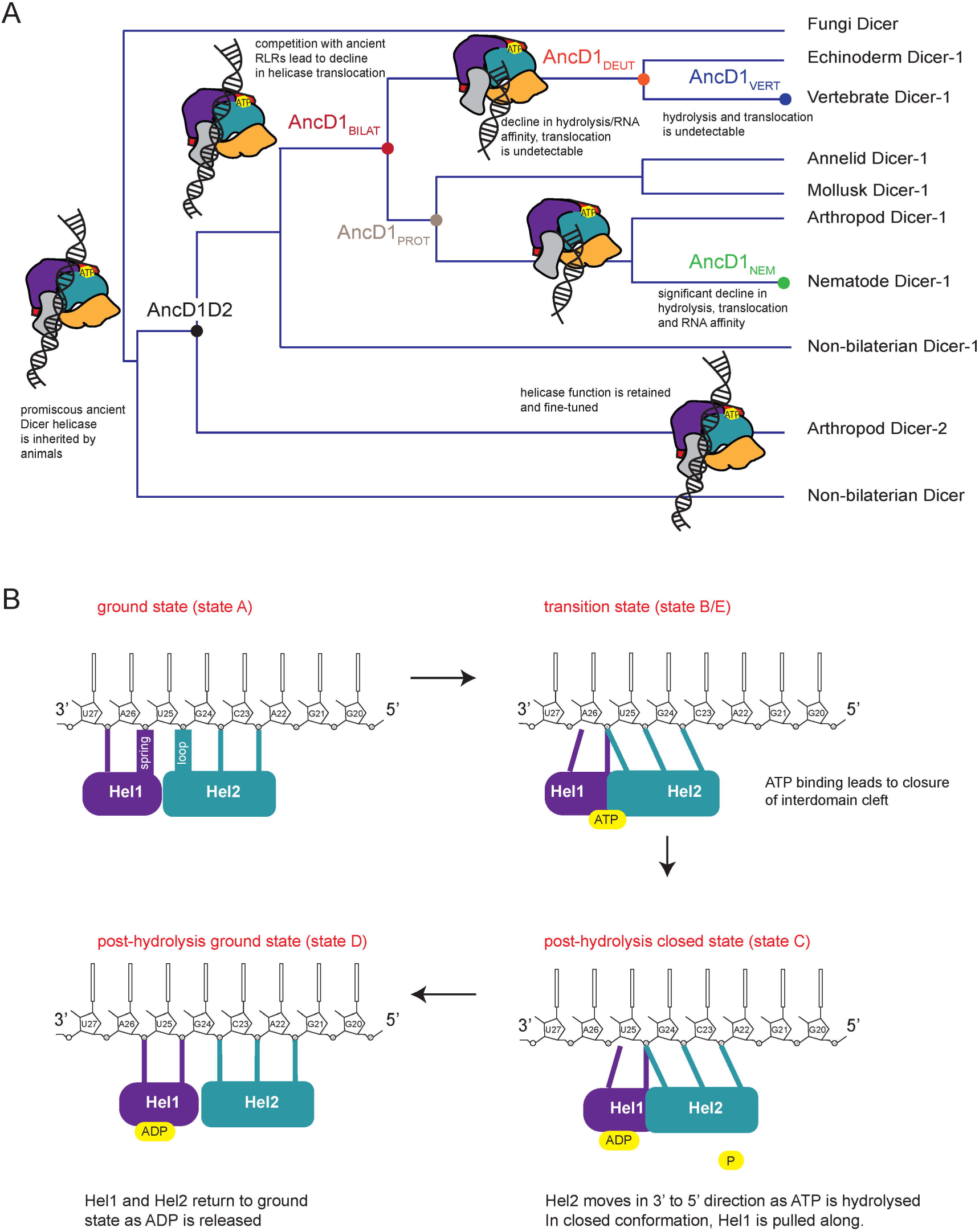
Models of metazoan Dicer helicase translocation and evolution. **(A)** Model of metazoan Dicer helicase evolution with text noting ancestral protein features. dsRNA at internal position signifies translocation. **(B)** Model for AncD1D2 translocation. Hel1 and Hel2, rounded rectangles. ATP binds ATPase pocket at interface of Hel1 and Hel2, causing Hel1 and Hel2 to move towards each another and form direct contacts. ATP hydrolysis leads to ejection of the γ-PO_4_ leading to conformational changes in Hel2 as motif VI loses contact with remaining ADP. Changes are propagated, causing Hel2 to loosen its grip on dsRNA and move 3’ to 5’, pulling Hel1 along. As ADP is released, AncD1D2-dsRNA is relaxed to a post hydrolytic ground state, disrupting interactions between Hel1 and Hel2, reestablishing the semi-closed state for another round of hydrolysis and 3’ to 5’ movement.

The flexible Hel2 loop provides an avenue for Hel2 to sample different phosphate backbone binding sites in the 5’ direction as evident in different loop conformations in D^post,ground^ subclasses (Movie S4). This flexibility may be directly tied to the 3’ to 5’ unidirectionality of RLR helicases and explain why mutation or deletion of this loop in RLRs eliminates translocation (23). Interestingly, Hel2 loop sidechains are observed to contact the 3’ strand phosphate backbone in RIG-I (38). Such a mechanism is likely generalizable between Dicer helicases and diverse RLRs that couple helicase movement along dsRNA to different mechanical outcomes. Moreover, the absence of AncD1_DEUT_ in internal dsRNA segments could be due to an inability to connect ATP hydrolysis to Hel2 movement in the 5’ direction causing AncD1_DEUT_ to remain stuck at the dsRNA terminus as it cycles through semi-closed and closed states. A similar mechanism, involving the Hel2 loop, was proposed to explain RIG-I translocation-throttling at viral dsRNA terminus (23).

Here we present a full ATPase cycle consisting of the following steps (Fig. 7B, Movie S5): Hel1 moving in the 5’ direction to close the ATPase pocket upon ATP binding, forming a high-energy, closed state; closed Hel1 and Hel2 sliding along the 3’ tracking strand as γ-PO_4_ is removed from bound ATP; Hel1 and Hel2 relaxing to a semi-closed, ground state at a +1 phosphate backbone position. A new ATP is then bound to restart the cycle all over again.

## Materials and Methods

Detailed descriptions of procedures are in SI Appendix, Materials and Methods. Phylogenetic analysis was performed as described (12). Ancestral Dicer helicases and modern full-length Dicers were expressed In Sf9 insect cells. RNAs were procured from IDT. Proteins were purified by affinity, ion exchange or heparin chromatography, and gel filtration (12). Gel shift and ATP hydrolysis assays were as described (12). Streptavidin-displacement assays and cryo-EM data collection, refinement and validation protocols are in SI Appendix, Materials and Methods and SI Appendix, Table S1.

### Data availability

Density maps and coordinate data for states A, B, C, D and E are deposited at the Electron Microscopy Data Bank (EMDB) and Protein Data Bank (RSCB PDB) under EMD accession codes EMD-48678, EMD-48691, EMD-48697, EMD-48708 and EMD-48710, and PDB accession codes 9MW6, 9MW7, 9MW8, 9MX3 and 9MX5, respectively. Ancestral sequences are in *SI Appendix*, Table S3.

## Acknowledgments

We thank Helen Donelick and Deirdre Mack for helpful discussion and training on cryo-EM, and members of the Bass Lab for helpful discussions and feedback. For cryo-EM work, we acknowledge David Belnap and Barbie Ganser-Pornillos at the University of Utah Electron Microscopy Core Laboratory. This work was supported by funding to B.L.B. (R35GM141262) and P.S.S. (R35GM133772) from the National Institute of General Medical Sciences of the NIH. B.L.B. is a Jon M. Huntsman Presidential Endowed Chair.

## Notes

### Competing Interest Statement

The authors have declared no competing interest.

## References

1. S. Paro, J.-L. Imler, C. Meignin, Sensing viral RNAs by Dicer/RIG-I like ATPases across species. Curr Opin Immunol 32, 106–113 (2015).

2. M. E. Fairman-Williams, U.-P. Guenther, E. Jankowsky, SF1 and SF2 helicases: family matters. Curr. Opin. Struct. Biol. 20, 313–324 (2010).

3. H. Kim, Y.-Y. Lee, V. N. Kim, The biogenesis and regulation of animal microRNAs. Nat. Rev. Mol. Cell Biol. 1–21 (2024). 10.1038/s41580-024-00805-0.

4. K. Jouravleva, et al., Structural basis of microRNA biogenesis by Dicer-1 and its partner protein Loqs-PB. Mol. Cell 82, 4049–4063.e6 (2022).

5. K. Förstemann, M. D. Horwich, L. Wee, Y. Tomari, P. D. Zamore, Drosophila microRNAs Are Sorted into Functionally Distinct Argonaute Complexes after Production by Dicer-1. Cell 130, 287–297 (2007).

6. N. K. Sinha, J. Iwasa, P. S. Shen, B. L. Bass, Dicer uses distinct modules for recognizing dsRNA termini. Science 359, 329–334 (2018).

7. M. Naganuma, H. Tadakuma, Y. Tomari, Single-molecule analysis of processive double-stranded RNA cleavage by Drosophila Dicer-2. Nat Commun 12, 4268 (2021).

8. R. K. Singh, et al., Transient kinetic studies of the antiviral Drosophila Dicer-2 reveal roles of ATP in self–nonself discrimination. Elife 10, e65810 (2021).

9. S. Su, et al., Structural insights into dsRNA processing by Drosophila Dicer-2–Loqs-PD. Nature 1–8 (2022). 10.1038/s41586-022-04911-x.

10. Z. Liu, et al., Cryo-EM Structure of Human Dicer and Its Complexes with a Pre-miRNA Substrate. Cell 173, 1191–1203.e12 (2018).

11. Y.-Y. Lee, H. Lee, H. Kim, V. N. Kim, S.-H. Roh, Structure of the human DICER–pre-miRNA complex in a dicing state. Nature 1–8 (2023). 10.1038/s41586-023-05723-3.

12. A. M. Aderounmu, P. J. Aruscavage, B. Kolaczkowski, B. L. Bass, Ancestral protein reconstruction reveals evolutionary events governing variation in Dicer helicase function. Elife 12, e85120 (2023).

13. P. V. Maillard, et al., Antiviral RNA Interference in Mammalian Cells. Science 342, 235–238 (2013).

14. E. Z. Poirier, et al., An isoform of Dicer protects mammalian stem cells against multiple RNA viruses. Science 373, 231–236 (2021).

15. G. J. Seo, et al., Reciprocal Inhibition between Intracellular Antiviral Signaling and the RNAi Machinery in Mammalian Cells. Cell Host Microbe 14, 435–445 (2013).

16. A. G. van der Veen, et al., The RIG-I-like receptor LGP2 inhibits Dicer-dependent processing of long double-stranded RNA and blocks RNA interference in mammalian cells. Embo J 37, e97479 (2018).

17. C. Gurung, et al., Dicer represses the interferon response and the double-stranded RNA-activated protein kinase pathway in mouse embryonic stem cells. J Biological Chem 296, 100264 (2021).

18. T. F. Duchaine, et al., Functional Proteomics Reveals the Biochemical Niche of C. elegans DCR-1 in Multiple Small-RNA-Mediated Pathways. Cell 124, 343–354 (2006).

19. A. Ashe, et al., A deletion polymorphism in the Caenorhabditis elegans RIG-I homolog disables viral RNA dicing and antiviral immunity. Elife 2, e00994 (2013).

20. C. D. Consalvo, et al., Caenorhabditis elegans Dicer acts with the RIG-I-like helicase DRH-1 and RDE-4 to cleave dsRNA. eLife 13, RP93979 (2024).

21. H. Jia, O. Kolaczkowski, J. Rolland, B. Kolaczkowski, Increased Affinity for RNA Targets Evolved Early in Animal and Plant Dicer Lineages through Different Structural Mechanisms. Mol Biol Evol 34, 3047–3063 (2017).

22. Z. Gao, M. Wang, D. Blair, Y. Zheng, Y. Dou, Phylogenetic Analysis of the Endoribonuclease Dicer Family. Plos One 9, e95350 (2014).

23. S. C. Devarkar, B. Schweibenz, C. Wang, J. Marcotrigiano, S. S. Patel, RIG-I Uses an ATPase-Powered Translocation-Throttling Mechanism for Kinetic Proofreading of RNAs and Oligomerization. Mol Cell 72, 355–368.e4 (2018).

24. K.-Y. Lee, C. Craig, S. S. Patel, Unraveling blunt-end RNA binding and ATPase-driven translocation activities of the RIG-I family helicase LGP2. Nucleic Acids Res. 52, 355–369 (2023).

25. A. Tsutsumi, T. Kawamata, N. Izumi, H. Seitz, Y. Tomari, Recognition of the pre-miRNA structure by Drosophila Dicer-1. Nat. Struct. Mol. Biol. 18, 1153–1158 (2011).

26. H. M. Donelick, et al., In vitro studies provide insight into effects of Dicer-2 helicase mutations in Drosophila melanogaster. Rna 26, 1847–1861 (2020).

27. C. D. Consalvo, et al., C. elegans Dicer acts with the RIG-I-like helicase DRH-1 and RDE-4 to cleave dsRNA. (2024). 10.7554/elife.93979.2.

28. D. C. Rawling, A. S. Kohlway, D. Luo, S. C. Ding, A. M. Pyle, The RIG-I ATPase core has evolved a functional requirement for allosteric stabilization by the Pincer domain. Nucleic Acids Res 42, 11601–11611 (2014).

29. M. Gu, C. M. Rice, Three conformational snapshots of the hepatitis C virus NS3 helicase reveal a ratchet translocation mechanism. Proc National Acad Sci 107, 521–528 (2010).

30. X. Ren, M. M. Linehan, A. Iwasaki, A. M. Pyle, RIG-I Selectively Discriminates against 5′-Monophosphate RNA. Cell Reports 26, 2019–2027.e4 (2019).

31. E. Uchikawa, et al., Structural Analysis of dsRNA Binding to Anti-viral Pattern Recognition Receptors LGP2 and MDA5. Mol Cell 62, 586–602 (2016).

32. Q. Yu, K. Qu, Y. Modis, Cryo-EM Structures of MDA5-dsRNA Filaments at Different Stages of ATP Hydrolysis. Mol Cell 72, 999–1012.e6 (2018).

33. E. Kowalinski, et al., Structural Basis for the Activation of Innate Immune Pattern-Recognition Receptor RIG-I by Viral RNA. Cell 147, 423–435 (2011).

34. X. Ye, L. Mayne, S. W. Englander, A conserved strategy for structure change and energy transduction in Hsp104 and other AAA+ protein motors. J. Biol. Chem. 297, 101066 (2021).

35. D. Zapletal, et al., Structural and functional basis of mammalian microRNA biogenesis by Dicer. Mol Cell 82, 4064–4079.e13 (2022).

