## Supplementary Material for "Biochemical and structural basis of Dicer helicase function unveiled by resurrecting ancient proteins"

#### Supplementary Materials and Methods

##### Phylogenetic tree construction and ancestral sequence reconstruction

Tree construction and prediction of ancestral sequences was as described (1). In addition, we manually constrained the Bilaterian Dicer-1 clade of the gene tree to a phylogeny resembling the consensus bilaterian species phylogeny. Tree editing was performed with Archaeopteryx software (https://github.com/cmzmasek/forester). The species-constrained tree was then rerun through RaxML-NG in evaluation mode to compute likelihood and transfer bootstrap values (2, 3). The species tree was used to predict amino acid sequences at select nodes as described (1).

##### Cloning, overexpression and purification of recombinant proteins

Production of ancestral and extant recombinant constructs was as described (1, 4, 5). DNA sequences were codon-optimized for expression in *Spodoptera frugiperda* (Sf9) insect cells using Integrated DNA Technologies' (IDT) codon optimization tool. cDNA sequences were synthesized by IDT with flanking primer sites for subcloning into a modified pFastBac plasmid containing 2×-Strep Flag tag, with NEB HiFi DNA Assembly Kit. Plasmid sequence was confirmed by sequencing (Plasmidsaurus) and transformed into Dh10Bac *E. coli* cells to generate bacmids, for transfection into Sf9 cells to produce baculovirus vectors for protein expression. After 2 rounds of baculovirus scale-up, "P2" baculovirus was used to infect large scale Sf9 cultures for protein production. Baculovirus titer was tracked after each virus amplification step using flow cytometry. Ancestral HEL-DUF constructs were purified using Strep-Tactin affinity chromatography, heparin chromatography or ion exchange chromatography, and size exclusion chromatography. Ancestral constructs eluted as monomers except $ANCD1_{VERT}$, which eluted as a mixture of monomers and dimers. Purified constructs were identified with LC/MS/MS at University of Utah Metabolomics Core. ceDCR-1 complexes and dmDcr2 were expressed and purified as described (4, 5).

##### RNA synthesis and sequence

RNA sequences were produced by chemical synthesis at IDT. Equimolar amounts of ssRNAs were annealed in annealing buffer (50 mM TRIS pH 8.0, 20 mM KCl) by placing the reaction on a heat block (95°C) and slow cooling ≥2 hrs (4). dsRNAs were gel purified after 8% polyacrylamide native PAGE and quantified using a Nanodrop or scintillation counter for 32P-labelled dsRNA.

42-nt sense ssRNA:
5'-GGGAAGCUCAGAAUAUUGCACAAGUAGAGCUUCUCGAUCCCC-3'

42-nt BLT-BLT antisense ssRNA:
5'-GGGGAUCGAGAAGCUCUACUUGUGCAAUAUUCUGAGCUUCCC-3'

42-nt 3'ovr-3'ovr antisense ssRNA:
5'-GGAUCGAGAAGCUCUACUUGUGCAAUAUUCUGAGCUUCCCGG-3'

40-nt sense 3'ovr-BLT ssRNA:
5'-GAAGCUCAGAAUAUUGCACAAGUAGAGCUUCUCG

42-nt sense deoxy ssRNA:
5'-GGGAAGCUCAGAAUAUUGCACA**dAdGdUdAdGdA**GCUUCUCGAUCCCC-3'

42-nt BLT-BLT antisense ssRNA:
5'-GGGGAUCGAGAAGCUCU**dAdCdUd**UGUGCAAUAUUCUGAGCUUCCC-3'

27-nt BLT-BLT sense ssRNA:
5'-AUACGUCCUGAUAGUUAGUAUCCAUCG-3'

27-nt BLT-BLT antisense ssRNA:
5'-CGAUGGAUACUAACUAUCAGGACGUAU-3'

**Thin-layer Chromatography ATP hydrolysis assay**

All reactions were performed at 37°C. Reaction mixtures containing 5X cleavage assay buffer (final concentrations: 25 mM TRIS pH 8.0, 100 mM KCl, 10 mM $MgCl_2$, 1 mM TCEP) was mixed with 5X ATP (500μM) and 5X [α-$^{32}$P] ATP (3000 Ci/mmol, 500nM). Reactions were started by mixing previous solution with 5X dsRNA (2μM) and 5X protein (1μM). Mixing equal volumes of all listed ingredients resulted in reaction with the following final conditions: 200nM protein, 400nM dsRNA, 100μM ATP and [α-$^{32}$P] ATP (3000 Ci/mmol, 100nM). Final reaction volume was incubated and aliquots of 2μL were removed at indicated times, quenched by addition of 3μL 500mM EDTA. 3μL of stopped reaction was spotted onto pre-washed 20×20 cm PEI-cellulose plates (Cel 300 PEI/UV 254 thin-layer chromatography (TLC) Plates 20x20, Machery-Nagel, Ref 801063), and chromatographed with 0.75 M $KH_2P0_4$ (adjusted to pH 3.3 with $H_3PO_4$) until solvent front reached the top of the plate. Plates were dried, visualized on a PhosphorImager screen (Molecular Dynamics), and quantified using ImageQuant version 8 software. Data was analyzed and visualized with GraphPad Prism Version 10.2.3.

Quantification of ATP hydrolysis assays for Table S1 was done by fitting the data into a two-phase exponent, with the first phase modeled as a linear reaction between time 0 and 2.5 min. The first phase is considered a transient zero order reaction where the rate constant $k_{(BURST)}$ is equal to the velocity of the reaction, which is the slope of ADP produced/ATP consumed (y) per minute (t). This rapid first phase probably ended before 2.5 min, but we are limited by the nature of manually mixed assays, as opposed to stopped flow assays where mixing and signal collection can be done on the timescale of seconds.

$y=k_{(BURST)}t + \text{intercept}$

Data for the second phase were fit to the pseudo-first order equation y=yo + A × (1-e−kt); where y=product formed (ADP in μM); A=amplitude of the rate curve, yo = baseline, k=pseudo-first-order rate constant = $k_{obs(TLC)}$; t=time. Data points are mean ± SD (n≥3).

**Colorimetric ATP hydrolysis assay**

These ATP hydrolysis reactions used the Quantichrom ATPase/GTPase Assay Kit (BioAssay Systems). Reactions were at 37°C in 108μL with cleavage assay buffer (final: 25mM TRIS pH 8.0, 100mM KCl, 10mM $MgCl_2$, 1mM TCEP), 2μM dsRNA and 0.05-2mM ATP; AncD1D2 was 25nM due to high ATP hydrolysis observed in TLC plate assays, but $AncD1_{BILAT}$ and $AncD1_{DEUT}$ were 100nM. Reaction mix of protein and dsRNA was preincubated at 37°C for 5 minutes before addition of ATP to start reaction. At timepoints, 18μL was removed and added to 2μL of 500mM EDTA to quench. As control, reaction mix of cleavage buffer and ATP was incubated alongside. 20μL reactions were plated on 96-well plates, and control was plated in duplicate. Phosphate standards were prepared as specified in assay kit, and 20uL of each were plated. After plating, 100μL of colorimetric reagent from the assay kit was added to each well, incubated for 30 minutes, and read on a Biotek Synergy Neo2 plate reader at 620nm. If necessary to fall within standard curve, reactions were diluted by four with water when plating. $OD_{620}$ values were fit to a linear regression, where velocity is equal to slope: ADP produced (μM) = velocity (μM/min) × time (min).

Velocities for each ATP concentration were fit to the Michaelis-Menten equation: $Y=Et*k_{cat}*X/(K_M + X)$, where Et is protein concentration (μM), $k_{cat}$ is turnover number, X is ATP concentration (mM), and $K_M$ is the Michaelis-Menten constant. Analysis used GraphPad Prism Version 10.2.3.

**Streptavidin-displacement assay**

Reactions were at 37°C in 100μL translocation assay buffer (final: 25mM HEPES, 10mM $Mg(OAc)_2$, 1mM TCEP). Reaction mixtures were prepared by incubating 1nM $^{32}$P labeled dsRNA with 2μM tetravalent streptavidin for 30 minutes. For reactions with ATP excluded, 0.010 units of hexokinase and 10mM glucose was added to deplete contaminating ATP. For reactions with ATP, 5mM ATP was added. Reactions were started by adding 1mM biotin and 200nM protein, except ceDCR-1 complexes which were 50nM protein. At indicated times, 12μL was removed and added to 3μL of 5X stop buffer (final: 25mM EDTA, 0.2% SDS, 10% glycerol, xylene cyanol, bromophenol blue). For AncD1D2, 2μM unlabeled trap dsRNA was added to stopping buffer to dislodge helicase-bound dsRNA and facilitate quantification of streptavidin-bound dsRNA. For ceDCR-1/DRH-1/RDE-4 complexes and dmDcr2$^{RIII}$, reactions were at 20°C and 25°C respectively. Stopping reaction mix was fractionated on 10% native 19:1 polyacrylamide gel, visualized on a PhosphorImager screen, and quantified with ImageQuant version 8.

Radioactivity in gels was quantified to determine fraction of streptavidin-bound and free dsRNA.
% streptavidin displaced = (free dsRNA/streptavidin-bound dsRNA) * 100
Rate constant, $k_{obs(STREP)}$, was determined by fitting data to a pseudo-first order rate equation $y=y_o + A \times (1-e^{-kt})$, where y is % streptavidin displaced; $y_o$ is baseline (~0); k is rate constant; t is time in min; A is amplitude.

**Electrophoretic mobility shift assay**

Electrophoretic mobility shift assays (EMSAs) were conducted with 500pM 42-basepair BLT dsRNA, with the 5' terminus of the sense strand labeled with $^{32}P$. Ancestral helicases were serially diluted in binding buffer (25mM TRIS pH 8.0, 100mM KCl, 10mM $MgCl_2$, 10% [vol/vol] glycerol, 1mM TCEP) to reach final concentrations of 0 – 1µM for AncD1D2, 0 – 5µM for $AncD1_{BILAT}$, 0 – 10µM for $AncD1_{DEUT}$. Labelled dsRNA was incubated with helicase dilutions in the presence of 5mM ATP-$MgOAc_2$ for 30 minutes at 37°C, and the reaction was stopped by loading directly onto a 5% native polyacrylamide gel (19:1 acrylamide/bisacrylamide) in 0.5X TRIS/Borate/EDTA running buffer. Gel was pre-run for 30 minutes before loading. Gels were electrophoresed (2 hr) at room temperature to resolve HEL-DUF bound to dsRNA from free dsRNA, dried (80°C, 1 hr) and exposed overnight to a PhosphorImager screen (Molecular Dynamics). dsRNA fraction that ran most slowly in the gel was attributed to bound dsRNA.

Radioactivity in gels corresponding to $dsRNA_{total}$ and $dsRNA_{free}$ was quantified using ImageQuant version 8 software to determine the fraction of dsRNA bound (Fraction bound = 1 – ($dsRNA_{free}/dsRNA_{total}$). To determine $K_D$ values, binding isotherms were fit using the Hill equation, where fraction bound = $1/(1 + [K_D^n/[P]^n])$; $K_D$ = equilibrium dissociation constant, n=Hill coefficient, and [P]=protein concentration. GraphPad Prism version 10.3.1 was used for curve-fitting analysis

**Cryo-EM sample preparation and data collection**

Samples were prepared by mixing AncD1D2 with 27-bp BLT dsRNA and incubating with appropriate nucleotide as described. For $A^{0ground}$ (AncD1D2 with dsRNA, no nucleotide), sample was prepared by mixing 6µM AncD1D2 with 7µM 27-bp BLT dsRNA and incubated on ice for ~30 minutes before application to grid. For $B^{end‡}$ (AncD1D2 with dsRNA, ADP-AlFx), 3µM AncD1D2 was incubated with 5µM 27-bp BLT dsRNA and a cocktail containing $MgCl_2$, $Al(NO_3)_3$, NaF and ATP at 37ºC for 30 minutes. For $C^{post,closed}$ and $D^{post,ground}$ (AncD1D2 with dsRNA, ATP), 6µM AncD1D2 was incubated with 15µM 27-bp BLT dsRNA and 2mM ATP at 37ºC for ~5 minutes and then transferred to ice for ~30 minutes before application to grid for blotting. For $E^{internal‡}$ (AncD1D2 with dsRNA, ADP-AlFx), 3µM AncD1D2 was incubated with 5µM 27-bp BLT dsRNA and a cocktail containing $MgCl_2$, $Al(NO_3)_3$, NaF and ATP at 37C for ~20 minutes and transferred to ice before application to grid. For $AncD1_{DEUT}$, 4.5µM was incubated with 15µM 27-bp BLT dsRNA and 2mM ATP at 37ºC for ~10 minutes and transferred to ice or incubated on ice overnight, before application to grid.

For AncD1D2 $C^{post,ground}$ and $D^{post,closed}$, Quantifoil R1.2/1.3 Cu300 mesh grids (SPT Labtech) were glow discharged for 100s at 25mA using a Pelco easiGlow unit (Ted Pella, Inc.) and 3µL of freshly prepared sample was applied to grids and blotted with filter paper (595 Filter Paper, Ted Pella, Inc.) for 7s using a Mk II Vitrobot (Thermo Fisher Scientific). For $AncD1_{DEUT}$ and AncD1D2 $A^{0ground}$, $B^{end‡}$ and $E^{internal‡}$, 3µL of freshly prepared sample was applied to glow discharged Quantifoil R1.2/1.3 Cu400 mesh grids and blotted with filter paper (595 Filter Paper, Ted Pella, Inc.) for 5s using a Mk IV Vitrobot (Thermo Fisher Scientific). Vitrobot conditions were set to 100% humidity, 4ºC temperature, 25 s wait time and -1mm offset. Grids were plunged into liquid ethane and frozen grids were transferred and stored in liquid nitrogen until data collection.

**CryoEM data acquisition**

CryoEM movies for all samples were recorded using EPU automated software (Thermo Fisher Scientific) on a 300 keV Titan Krios transmission electron microscope (Thermo Fisher Scientific) equipped with a post-GIF K3 direct detector (Gatan, Inc). State A (no nucleotide) and states C-E samples were collected at super-resolution with a nominal magnification of 105,000 x corresponding to a calibrated pixel size of 0.436 Å/pix. State B samples were collected at a nominal magnification of 105,000 x in counting mode, corresponding to a calibrated pixel size of 0.872 Å/pix. All samples were imaged with a defocus range of -0.8 μm to -2.4 μm at a total dose of 40 electrons/$Å^2$ and 50 frames per movie.

**Cryo-EM data processing**

All datasets were processed through cryoSPARC v4 (6). CryoEM movie frames were dose-weighted and beam-induced motion corrected (super-resolution movies were Fourier binned 2X). CTF parameter estimation was performed with Patch CTF Estimation. Motion-corrected micrographs were based on CTF fit resolution with a 5Å cutoff and manual inspection.

For state $A^{0ground}$, a total of 12,548,917 particles were selected from 11,003 micrographs, with a blob picker with particle diameter restraints of 60-120 Å. Particles were initially extracted with a box size of 320 pixels and Fourier cropped 4X to 80 pixels. After multiple rounds of 2D classification to discard junk and low-resolution particles, 951,277 particles were re-extracted at a box size of 320 pixels and used to generate 3D volumes with Ab initio 3D reconstruction. Volumes corresponding to a singlet and doublet helicase on 27-bp BLT dsRNA were used to perform heterogeneous reconstructions. 260,444 particles sorted into the doublet helicase volume were used as input for Topaz (7) to identify more particles in this doublet state. Particles from the other heterogenous refinement classes were also independently used to train Topaz. Newly extracted particles from Topaz were combined to run another round of heterogeneous refinement which yielded a doublet helicase class containing 303,540 particles. Non-uniform refinement followed by local refinement was then performed with a mask around the higher resolution helicase in the doublet resulting in a 3.4 Å map.

For state $B^{end‡}$, a total of 23,429,138 particles were selected from 11,834 micrographs, with a blob picker with particle diameter restraints of 60-120 Å. Particles were initially extracted with a box size of 288 pixels and Fourier cropped 4X to 72 pixels. After multiple rounds of 2D classification to discard junk and low-resolution particles, 435,015 particles were re-extracted at a box size of 288 pixels and used to generate 3D volumes with Ab initio 3D reconstruction. Two volumes corresponding to a singlet helicase on 27-bp BLT dsRNA were used to perform heterogenous reconstructions. 240,370 particles were sorted into a class with intact helicase features bound to the end of dsRNA. This particle set was used to train Topaz and the reextracted particles were subject to a second round of heterogeneous refinement. The resulting 380,191 particles were used as input for non-uniform refinement followed by local refinement around the helicase domain to generate a density map with 3.4 Å global resolution at 0.143 FSC threshold.

For states $C^{post,closed}$ and $D^{post,ground}$, a total of 21,532,216 particles were picked from 15,138 micrographs, with a blob picker with particle diameter restraints of 60-120 Å. Particles were extracted with a 288-pixel box size, Fourier-binned 4X to 72 pixels and subjected to multiple rounds of 2D classification. The resulting 2,256,542 particles were reextracted at a box size of 288 pixels and used to generate eight heterogeneous refinement classes after ab initio reconstruction. 4 of the classes contained intact AncD1D2 helicases bound to dsRNA while the remaining particles contained AncD1D2 bound to dsRNA but missing Hel2i, likely due to flexibility of Hel2i as observed with the state $A^{0ground}$ and $B^{end‡}$ structures. The intact AncD1D2 particles were subjected to focused 3D classification resulting in 4 classes, 2 of which were combined to form the consensus state D density map after non uniform refinement and local refinement. Of the remaining two classes, state C and state D.3 were generated after non uniform and local refinement. AncD1D2 particles missing Hel2i were also subject to 3D refinement and a single class (state D.4) was subject to non-uniform refinement followed by local refinement to generate a high-quality density map with a global resolution of 3.2 Å at 0.143 FSC threshold.

For state $E^{internal‡}$, a total of 12,806,533 particles were selected from 7,335 micrographs, with a blob picker with particle diameter restraints of 60-120 Å. Particles were initially extracted with a box size of 320 pixels and Fourier cropped 4X to 80 pixels. After multiple rounds of 2D classification to discard junk and low-resolution particles, 639,691 particles were re-extracted at a box size of 320 pixels and used to generate 3D volumes with Ab initio 3D reconstruction. Volumes corresponding to a singlet and doublet helicase on 27-bp BLT dsRNA were used to perform heterogeneous reconstructions. 261,953 particles sorted into the doublet helicase volume were used as input for Topaz to identify more particles in this doublet state. Particles from the other heterogeneous refinement classes were also independently used to train Topaz. Newly extracted particles from Topaz were combined to run another round of heterogeneous refinement which yielded a doublet helicase class containing 171,095 particles. Non-uniform refinement followed by local refinement was then performed with a mask around the higher resolution helicase in the doublet resulting in a 3.1 Å map.

**Model building refinement and validation**

The AlphaFold2 predicted model of AncD1D2 (Table S3) was used as starting point for initial model building followed by manual adjustment in Chimera, ChimeraX and COOT (8–11). ADP, Mg and AlFx were built and refined in PHENIX using the Ligand Fit tool (12). RNA nucleotides were fit manually using DeepEMhancer sharpened maps (13). All models were refined and validated against their respective density maps in PHENIX using the real space refinement tool and Molprobity (14, 15).

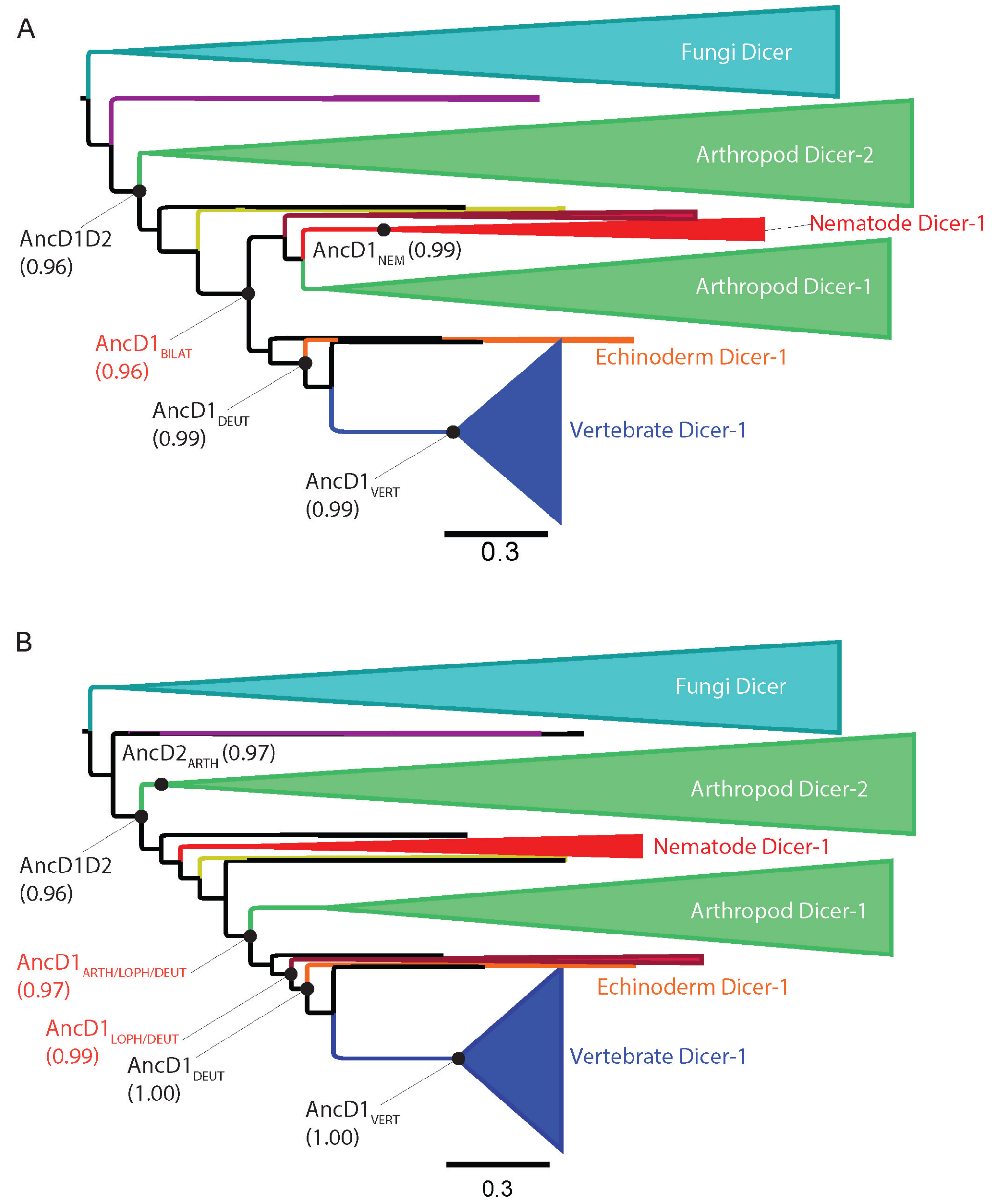


**Fig. S1. Phylogenetic trees constructed from extant metazoan Dicer HEL-DUF amino acid sequences. (A)** Dicer HEL-DUF phylogenetic tree with Dicer-1 clade constrained to species relationships, visualized and annotated with FigTree (http://tree.bio.ed.ac.uk/software/figtree/). Resurrected ancestral nodes are depicted with black circles with transfer bootstrap values indicated in parentheses. Width of cartooned triangle bases represents species sampling depth, i.e. number of species. Scale bar represents total amino acid substitutions divided by the number of amino acids, i.e., amino acid substitutions per site. Species tree-gene tree incongruent nodes are depicted with red labels. **(B)** Maximum likelihood Dicer HEL-DUF phylogenetic gene tree, visualized and annotated with FigTree. Resurrected ancestral nodes are depicted with black circles with transfer bootstrap values indicated in (parentheses). Width of cartooned triangle bases represents species sampling depth, i.e. number of species. Scale bar represents total amino acid substitutions divided by number of amino acid sites, i.e., amino acid substitutions per site. Species tree-gene tree incongruent nodes are depicted with red labels.

**42 BLT-BLT**

```
5'OH—GGGAAGCUCAGAAUAUUGCACAAGUAGAGCUUCUCGAUCCCC— 3'OH
3'OH—CCCUUCGAGUCUUAUAACGUGUUCAUCUCGAAGAGCUAGGGG— 5'OH
```

**42 3'ovr-3'ovr**

```
5'OH —GGGAAGCUCAGAAUAUUGCACAAGUAGAGCUUCUCGAUCCCC—3'OH
3'OH—GGCCCUUCGAGUCUUAUAACGUGUUCAUCUCGAAGAGCUAGG—5'OH
```

**42 BLT-BLT 5'p**

```
5'p —GGGAAGCUCAGAAUAUUGCACAAGUAGAGCUUCUCGAUCCCC— 3'OH
3'OH—CCCUUCGAGUCUUAUAACGUGUUCAUCUCGAAGAGCUAGGGG— 5'BIO
```

**42 3'ovr-BLT 5'p**

```
5'p — GAAGCUCAGAAUAUUGCACAAGUAGAGCUUCUCGAUCCCC—3'OH
3'OH—GGCCCUUCGAGUCUUAUAACGUGUUCAUCUCGAAGAGCUAGG—5'BIO
```

**42 BLT-BLT deoxy 5'p**

```
5'p —GGGAAGCUCAGAAUAUUGCACAAGUAGAGCUUCUCGAUCCCC— 3'OH
3'OH—CCCUUCGAGUCUUAUAACGUGUUCAUCUCGAAGAGCUAGGGG— 5'BIO
```

**27 BLT-BLT**

```
5'OH —AUACGUCCUGAUAGUUAGUAUCCAUCG— 3'OH
3'OH —UAUGCAGGACUAUCAAUCAUAGGUAGC— 5'OH
```

**Fig. S2. RNA duplexes used in this study.** 5' and 3' ends are labeled. Deoxynucleotides shown in red font. BIO is covalently linked biotin.

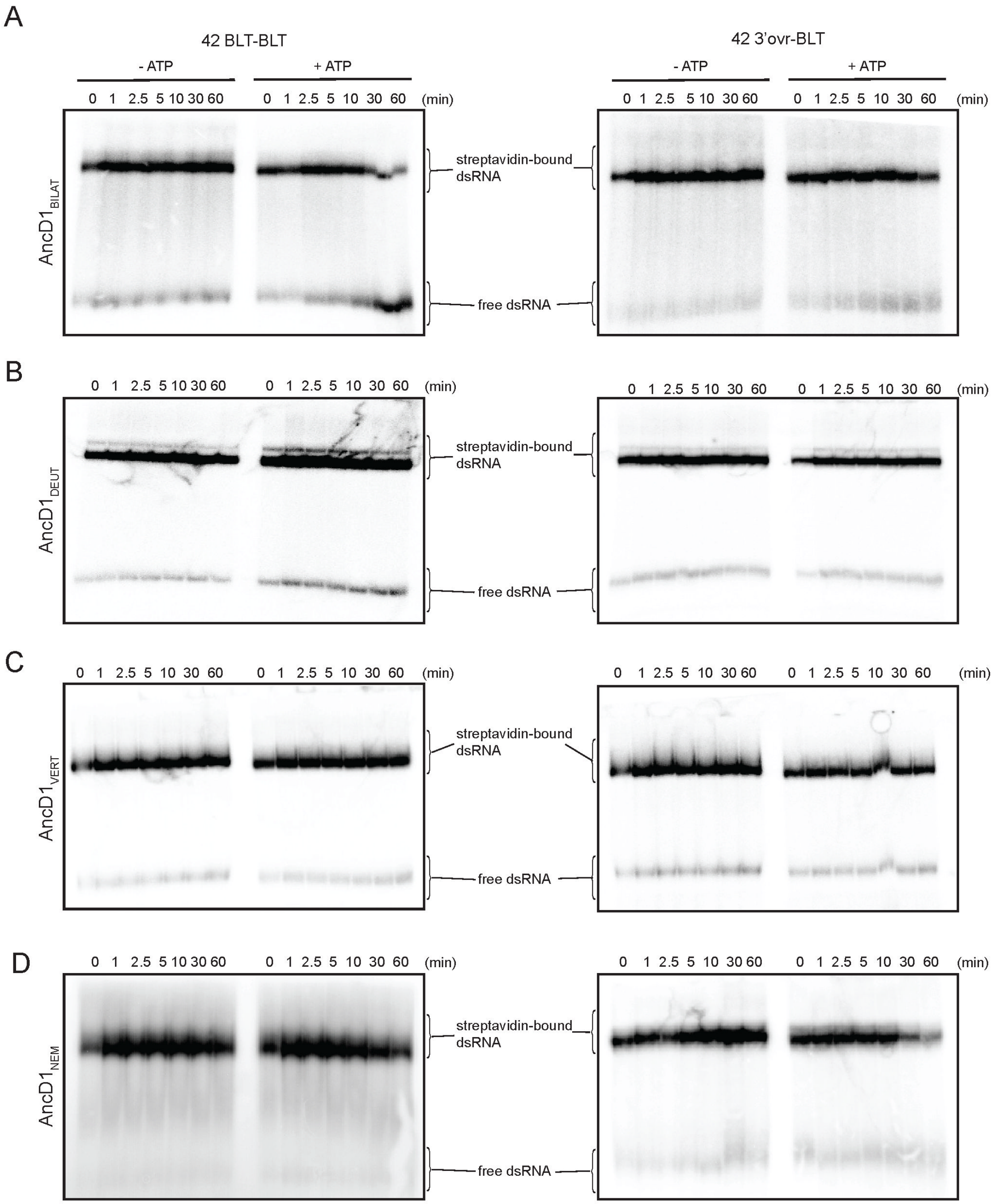


**Fig. S3. Ancestral Dicer helicases translocate on dsRNA. (A-D)** Representative PhosphorImages of streptavidin-displacement assays for **(A)** AncD1$_{BILAT}$ **(B)** AncD1$_{DEUT}$ **(C)** AncD1$_{VERT}$ and **(D)** AncD1$_{NEM}$, measuring translocation along BLT (left) and 3'ovr (right) dsRNA in the absence and presence of 5mM ATP.

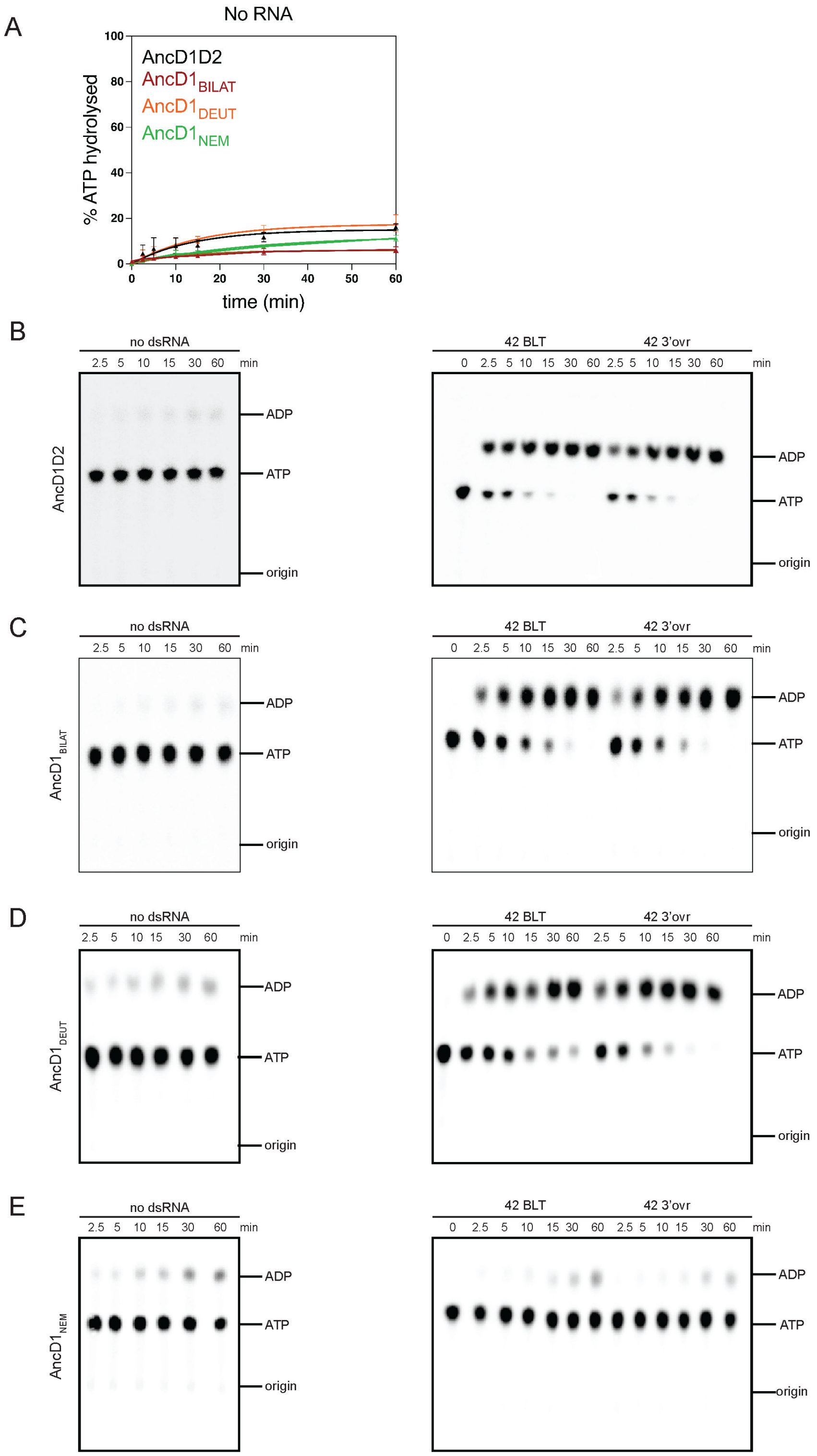


**Fig. S4. Ancient Dicer helicases hydrolyze ATP. (A)** Graph shows quantification of TLC analyses monitoring hydrolysis of ATP (100µM) by selected ancestral Dicer helicases in the absence of dsRNA. Data points are mean ± SD (n ≥ 3) and were fit to a pseudo-first order equation. **(B–E)** PhosphorImages of representative TLC plates showing hydrolysis of ATP (100µM spiked with α-$^{32}$P-ATP) by 200nM ancestral HEL-DUFs for various times as indicated, at 37°C, in the absence of dsRNA (left) or in the presence of 400nM 42-bp dsRNA with BLT or 3'ovr termini (right).

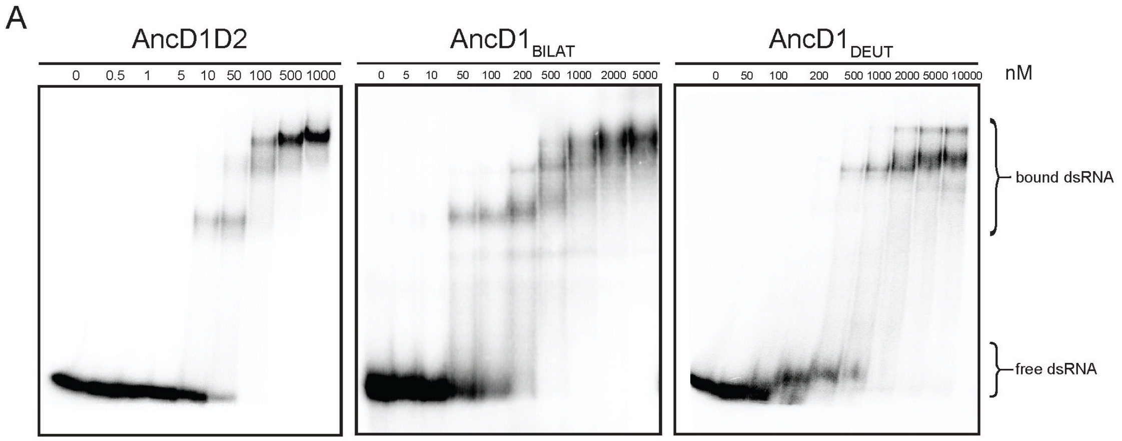


**Fig S5. Ancient Dicer helicases bind dsRNA.** Representative PhosphorImages of electrophoretic mobility shift assays for AncD1D2, $AncD1_{BILAT}$ and $AncD1_{DEUT}$.

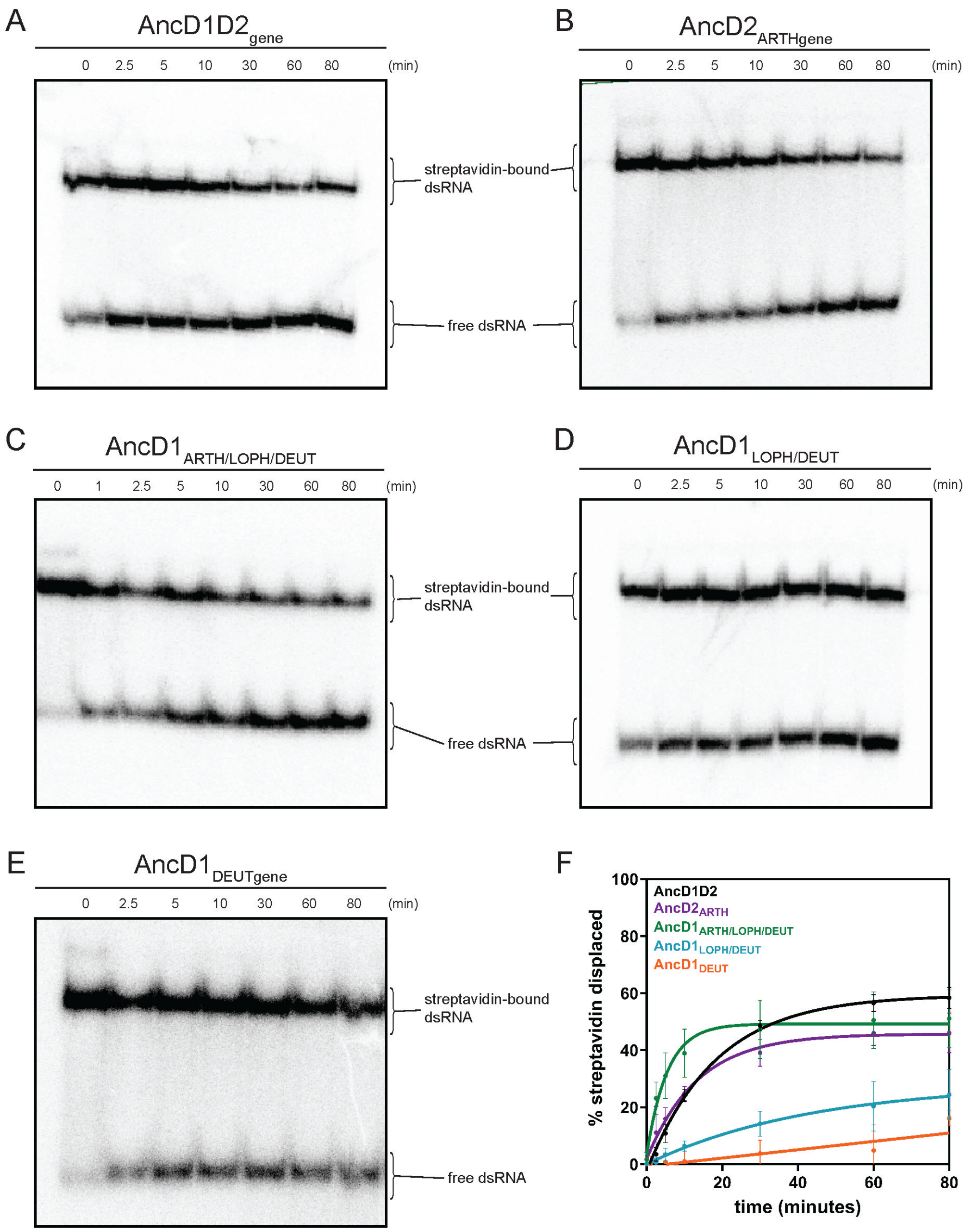


**Fig. S6. Ancient Dicer helicases from maximum likelihood tree translocate on dsRNA. (A-E)** Representative PhosphorImages of streptavidin-displacement assays for **(A)** AncD1D2 **(B)** AncD2$_{ARTH}$ **(C)** AncD1$_{ARTH/LOPH/DEUT}$ **(D)** AncD1$_{LOPH/DEUT}$ and **(E)** AncD1$_{DEUT}$ measuring translocation along BLT 42-bp dsRNA in the presence of 5mM ATP. **(F)** Graph showing streptavidin-displacement data for ancestral Dicer constructs from the maximum likelihood tree for BLT dsRNA. Data were fit to a pseudo-first order rate equation. Data points are mean ± SD (n ≥ 3).

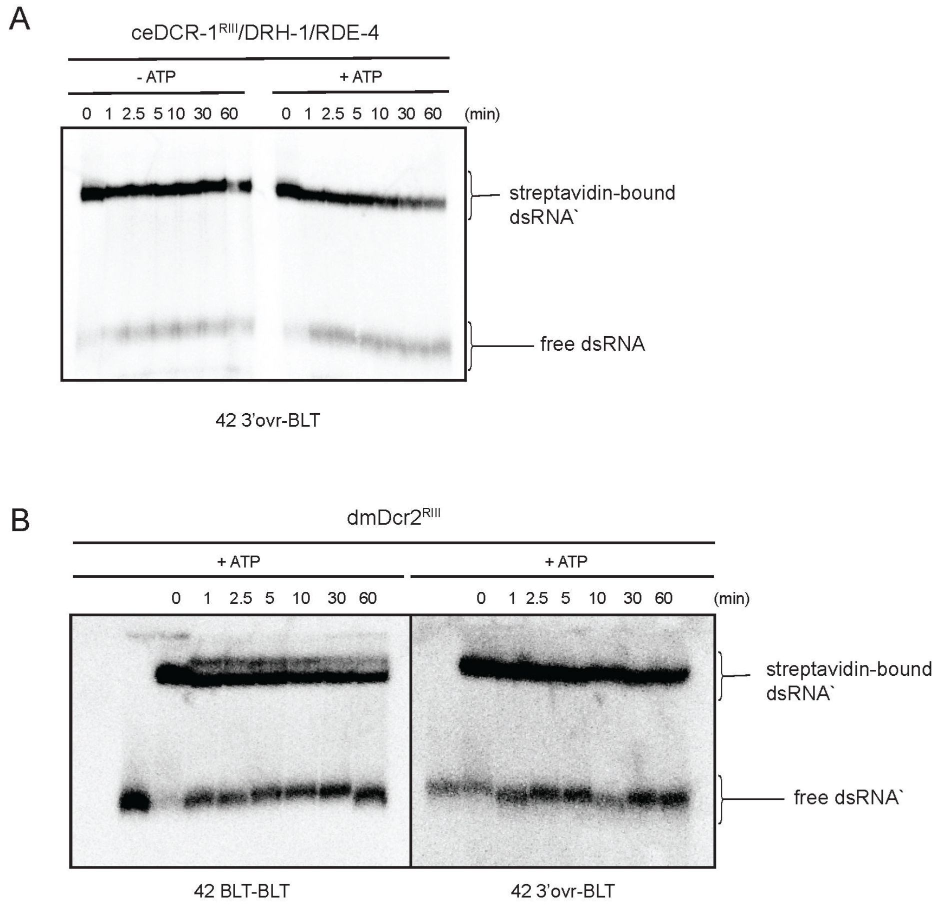


**Fig. S7. Invertebrate Dicers translocate along dsRNA. (A)** Representative PhosphorImage of streptavidin-displacement assay for ceDCR-1$^{RIII}$/DRH-1/RDE-4 translocation along 42-bp 3'ovr dsRNA in the absence and presence of 5mM ATP. **(B)** Representative PhosphorImage of streptavidin-displacement assay for dmDcr2$^{RIII}$ translocation along 42-bp BLT and 3'ovr dsRNA in the presence of 5mM ATP.

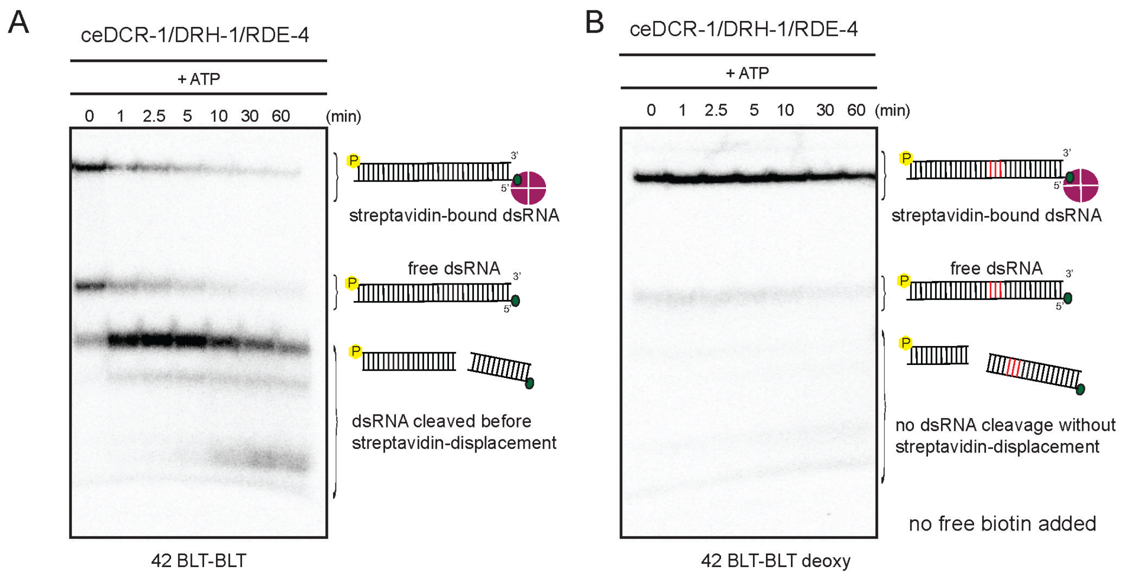


**Fig. S8. Streptavidin-displacement assay design shows that ceDCR-1 antiviral complex translocates along dsRNA. (A)** Representative PhosphorImage of streptavidin-displacement assay for ceDCR-1/DRH-1/RDE-4 translocation along BLT 42-bp dsRNA in the presence of 5mM ATP. dsRNA is cleaved instantly before translocation can be tracked. This showed that regular 42-bp dsRNA cannot be used to assay translocation of a wildtype Dicer as cleavage from the left side will precede streptavidin displacement from the right side. Cartoons indicate gel migration of different species. **(B)** Representative PhosphorImage of streptavidin-displacement assay for ceDCR-1/DRH-1/RDE-4 translocation along BLT 42-bp dsRNA with deoxynucleotides in the presence of 5mM ATP without free biotin sink for displaced streptavidin. Displaced streptavidin rapidly rebinds dsRNA and ceDCR-1 is unable to cleave dsRNA. This demonstrates that the deoxy patch effectively blocks cleavage from the left side. This assay also shows that streptavidin displacement is required for cleavage from the right side, which only occurs after free dsRNA reengages with the Dicer complex. Cartoons indicate deoxynucleotides (red) and gel migration of different species.

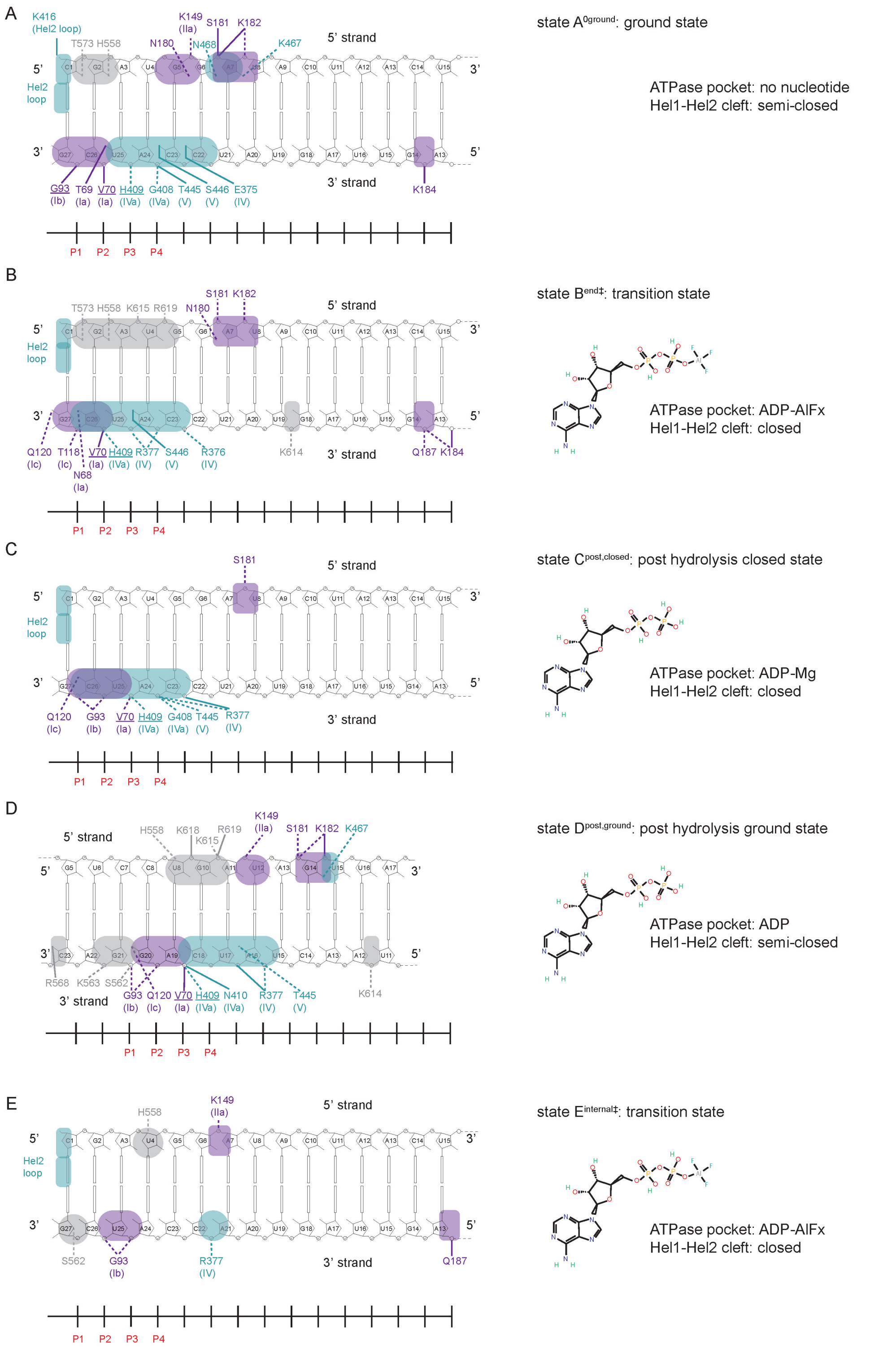


**Fig. S9. Schematic representation of AncD1D2-dsRNA interactions. (A)** state $A^{0ground}$ **(B)** state $B^{end‡}$ **(C)** state $C^{post,closed}$ **(D)** state $D^{post,ground}$ and **(E)** state $E^{internal‡}$. Key protein residues and motifs are highlighted, and sites of interaction are indicated with colored boxes. Residues used to monitor helicase movement along 3' tracking strand in Fig. 5 are underlined. Bold lines depict hydrogen bonds while dashed lines depict high confidence polar contacts. Nucleotide state is depicted with chemical structure showing position of aluminum fluoride in ATP mimics.

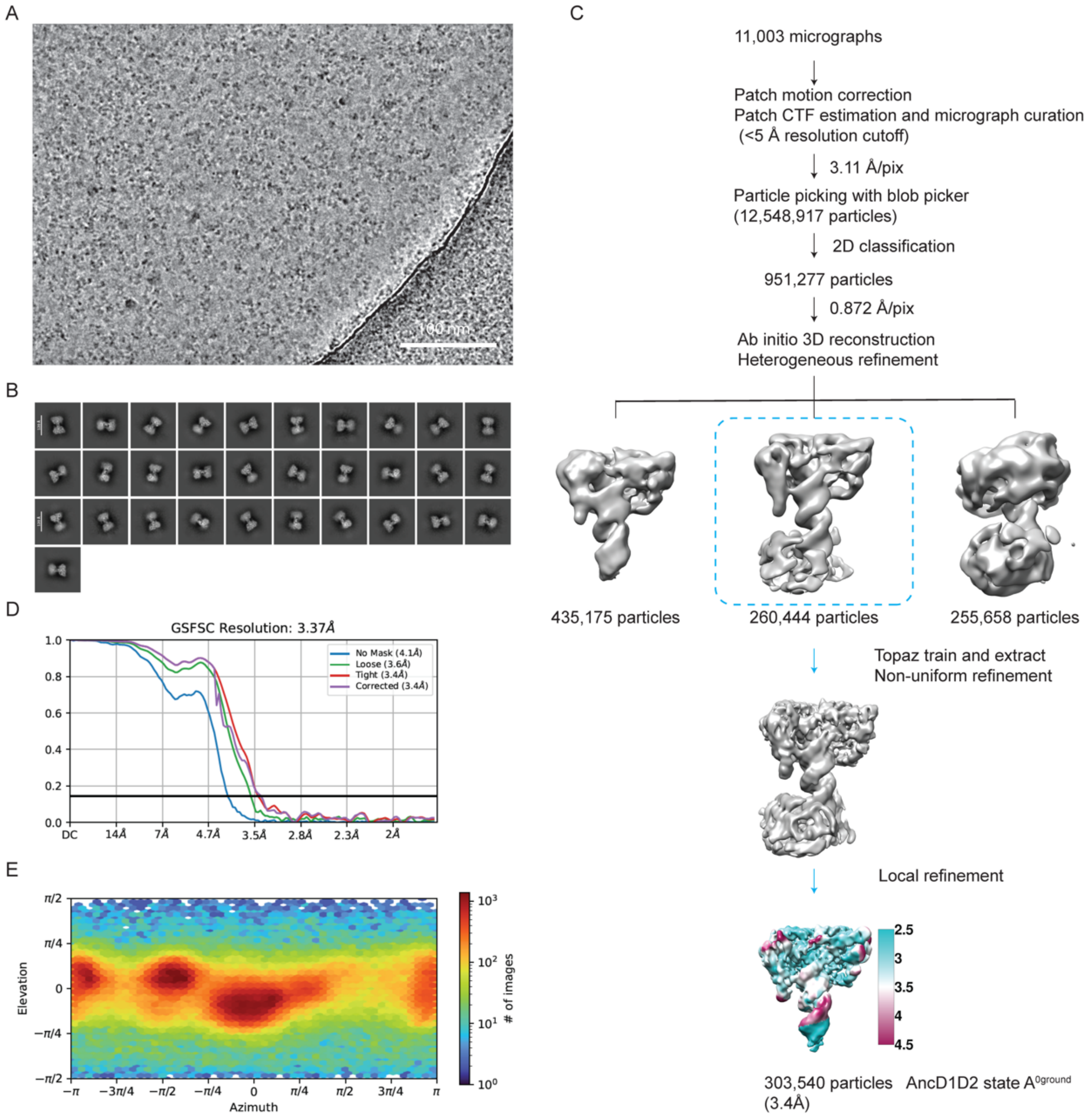


**Fig. S10. CryoEM image processing workflow of AncD1D2 in state $A^{0ground}$. (A)** Representative cryoEM micrograph of AncD1D2 bound to 27-bp dsRNA in state $A^{0ground}$**. (B)** Representative views of 2D class averages of AncD1D2 state $A^{0ground}$. **(C)** Flowchart of cryoEM data processing of the AncD1D2 state $A^{0ground}$. Colored map depicts local resolution. **(D)** Gold standard FSC of the final map of AncD1D2 state $A^{0ground}$. **(E)** Particle distribution orientation.

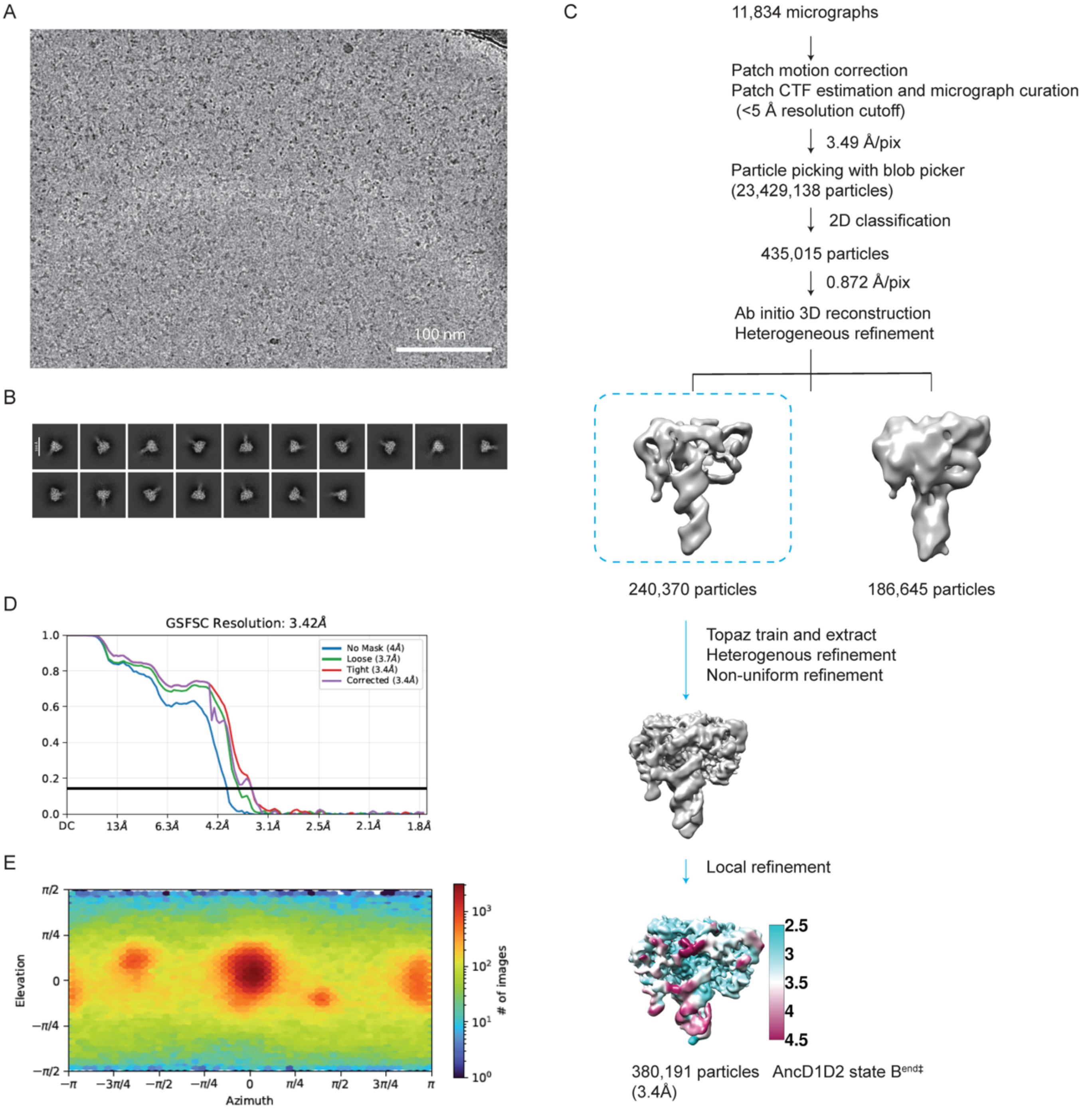


**Fig. S11. CryoEM image processing workflow of AncD1D2 in state B$^{end‡}$. (A)** Representative cryoEM micrograph of AncD1D2 bound to 27-bp dsRNA in state B$^{end‡}$. **(B)** Representative views of 2D class averages of AncD1D2 state B$^{end‡}$. **(C)** Flowchart of cryoEM data processing of the AncD1D2 state B$^{end‡}$. **(D)** Gold standard FSC of the final map of AncD1D2 state B$^{end‡}$. Colored map depicts local resolution. **(E)** Particle distribution orientation.

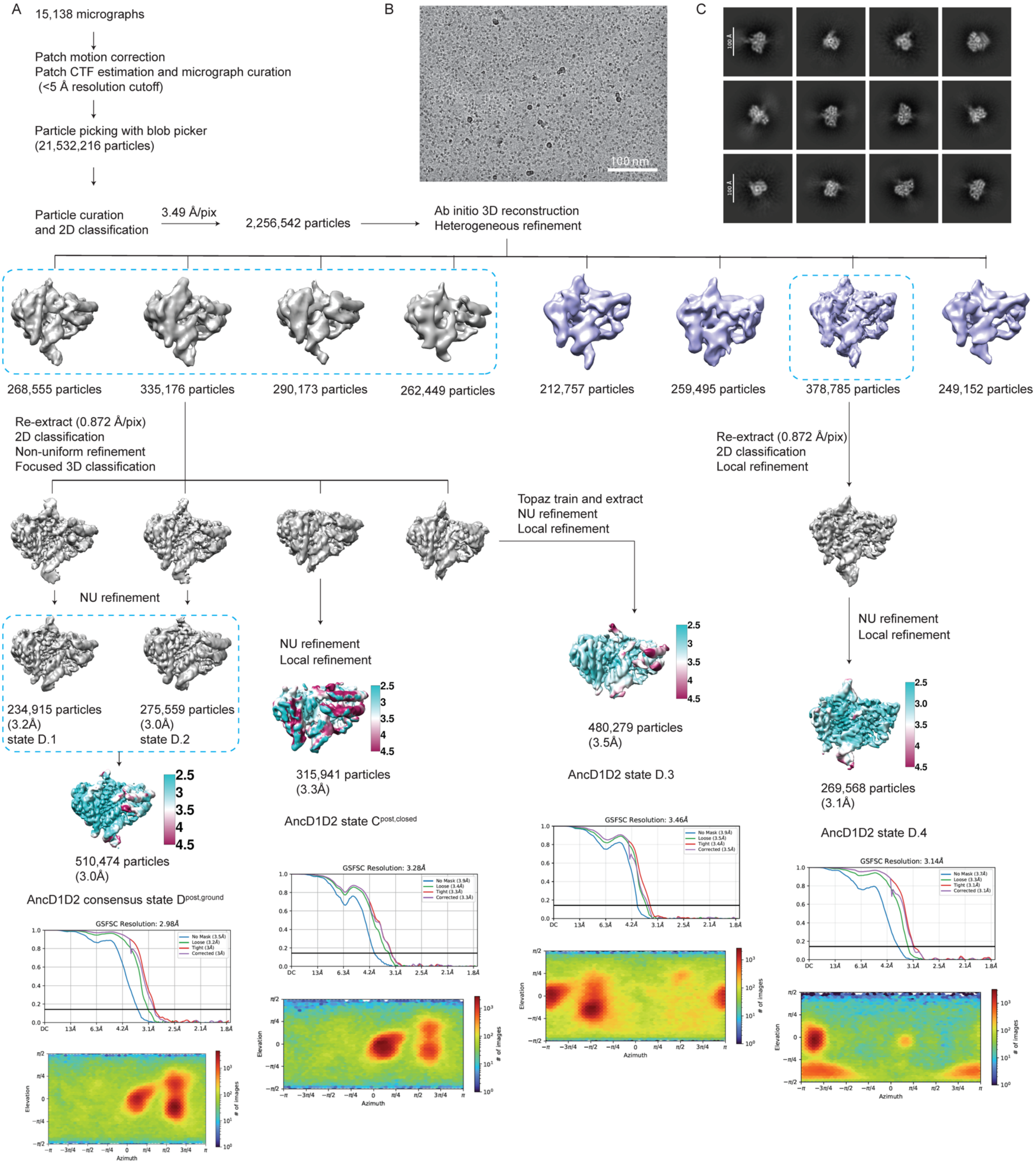
A
15,138 micrographs
Patch motion correction
Patch CTF estimation and micrograph curation
(<5 Å resolution cutoff)
Particle picking with blob picker
(21,532,216 particles)
Particle curation
and 2D classification
3.49 Å/pix
2,256,542 particles
Ab initio 3D reconstruction
Heterogeneous refinement
B
100 nm
C
100 Å
268,555 particles
335,176 particles
290,173 particles
262,449 particles
212,757 particles
259,495 particles
378,785 particles
249,152 particles
Re-extract (0.872 Å/pix)
2D classification
Non-uniform refinement
Focused 3D classification
Re-extract (0.872 Å/pix)
2D classification
Local refinement
Topaz train and extract
NU refinement
Local refinement
NU refinement
234,915 particles
(3.2Å)
state D.1
275,559 particles
(3.0Å)
state D.2
NU refinement
Local refinement
NU refinement
Local refinement
510,474 particles
(3.0Å)
AncD1D2 consensus state Dpost,ground
315,941 particles
(3.3Å)
AncD1D2 state Cpost,closed
480,279 particles
(3.5Å)
AncD1D2 state D.3
269,568 particles
(3.1Å)
AncD1D2 state D.4
GSFSC Resolution: 2.98Å
GSFSC Resolution: 3.28Å
GSFSC Resolution: 3.46Å
GSFSC Resolution: 3.14Å
Azimuth
Elevation
### of images

**Fig. S12. CryoEM image processing workflow of AncD1D2 in post hydrolysis states $C^{post,closed}$ and $D^{post,ground}$. (A)** Flowchart of cryoEM data processing of AncD1D2 bound to 27-bp dsRNA in post hydrolysis states $C^{post,closed}$ and $D^{post,ground}$. Colored map depicts local resolution. **(B)** Representative cryoEM micrograph of AncD1D2 bound to 27-bp dsRNA in post hydrolysis states $C^{post,closed}$ and $D^{post,ground}$. **(C)** Representative views of 2D class averages of AncD1D2 in post hydrolysis states $C^{post,closed}$ and $D^{post,ground}$.

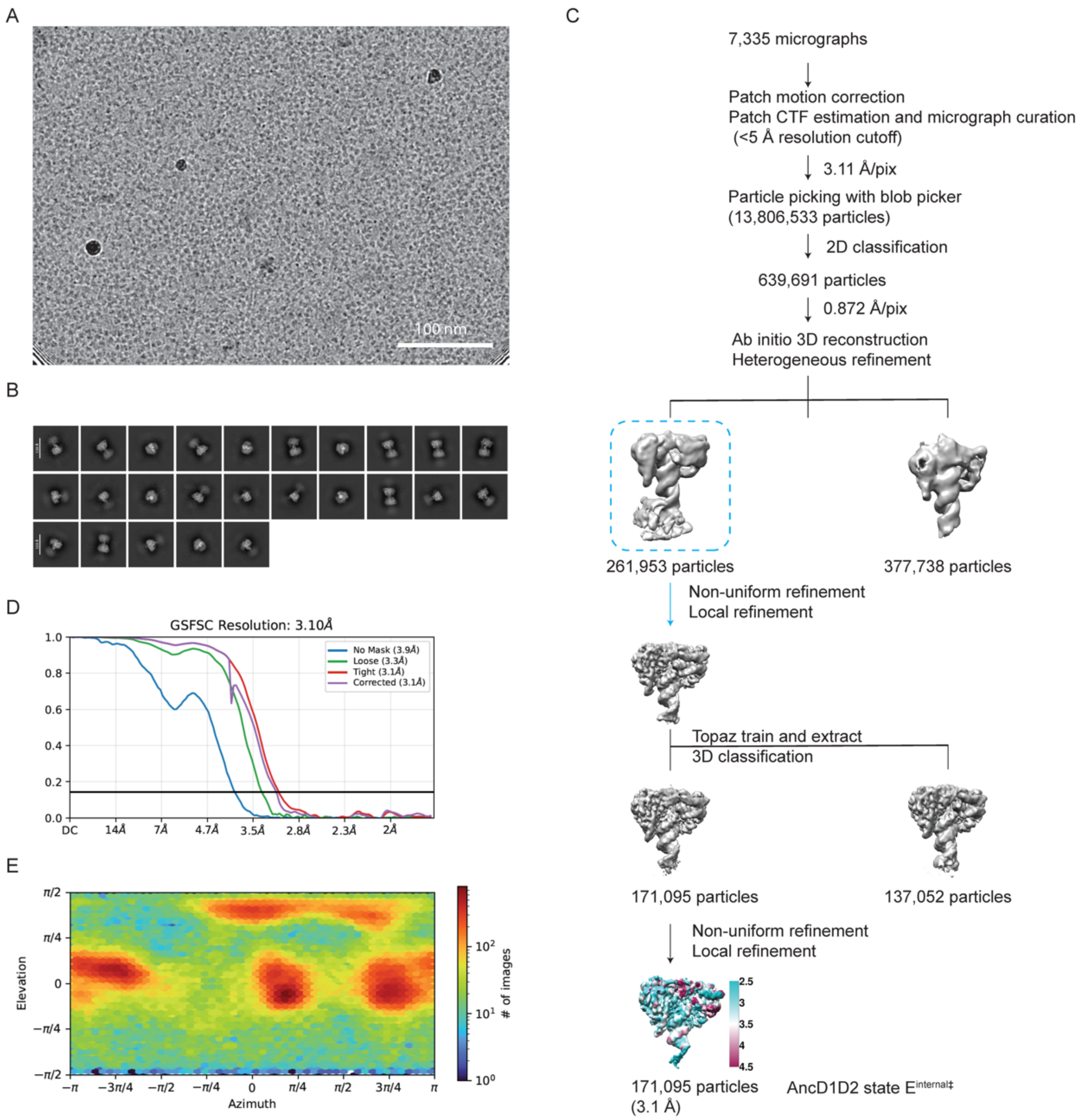


**Fig. S13. CryoEM image processing workflow of AncD1D2 in state $E^{internal‡}$. (A)** Representative cryoEM micrograph of AncD1D2 bound to 27-bp dsRNA in state $E^{internal‡}$. **(B)** Representative views of 2D class averages of AncD1D2 internal state $E^{internal‡}$. **(C)** Flowchart of cryoEM data processing of the AncD1D2 internal state $E^{internal‡}$. Colored map depicts local resolution. **(D)** Gold standard FSC of the final map of AncD1D2 internal state $E^{internal‡}$. **(E)** Particle distribution orientation.

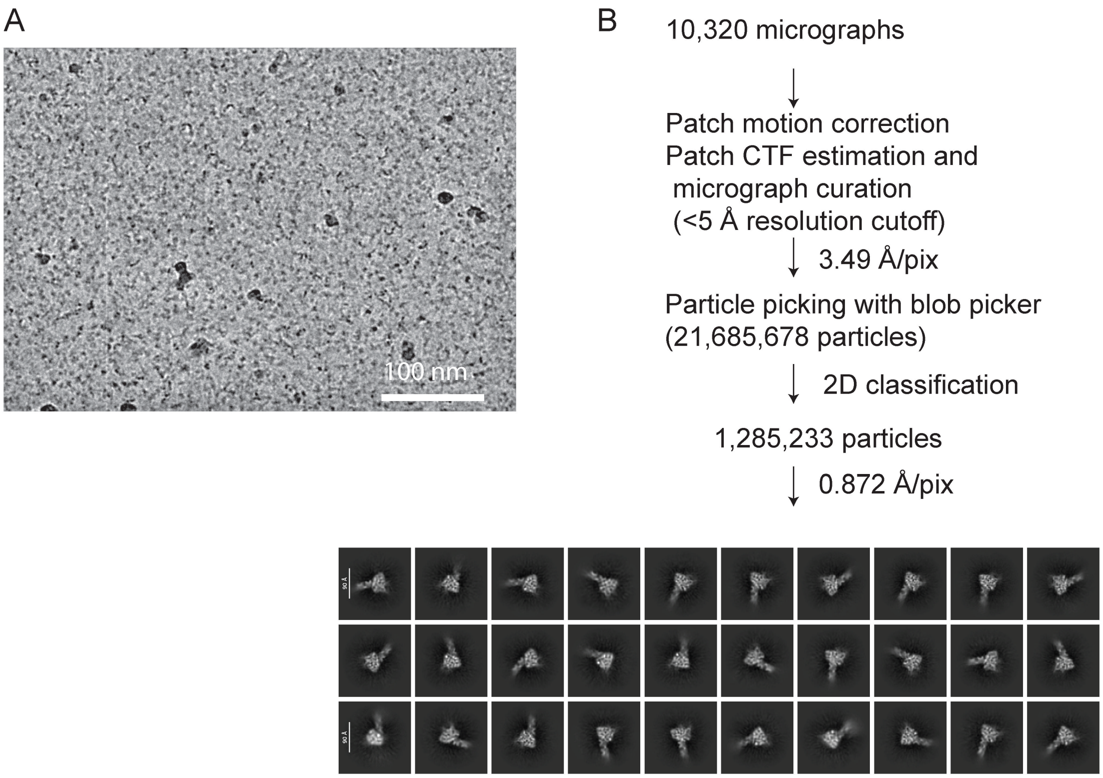


**Fig. S14. CryoEM image processing workflow of AncD1$_{\text{DEUT}}$ in endbound state. (A)** Representative cryoEM micrograph of AncD1$_{\text{DEUT}}$ bound to 27-bp BLT dsRNA in endbound state with added ATP. **(B)** Flowchart of cryoEM data processing of the AncD1$_{\text{DEUT}}$ in endbound state with added ATP. Processing terminates with 2D classification.

A

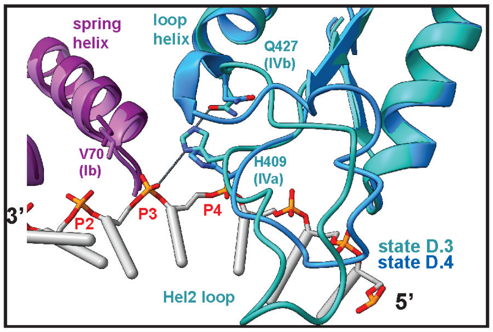


B

```
                                  H409                Q427
DMDCR2                 414  ELRNVLTPQFMVGR...NNISPDFESVLERKWQKSAIQQFRDGNA
ANCD2ARTH              399  D.FSFIKPDFIVGHNS.SPSSKEKSTEMSSKKQKEVLKKFRKGEC
ANCD1D2                399  D.FSFIKCDFIVGHNS.SPSSKEKSTEMSSKKQKEVLKKFRKGEL
ANCD1D2SPECIES         397  D.FSFIKCDFVVGHNS.SPSSKEKSTEMNSKKQKEVLKKFRKGEC
ANCD1ARTHLOPHDEUT      424  D.FSFIKSDYIVGHGTGSTGTSSKETEMQFRKQEEVLRKFRKHEC
ANCD1BILATSPECIES      429  D.FSFIKSDYIVGHGSGSTGTSSKETEMQFKKQEEVLRKFRRHEC
ANCD1NEMATODESPECIES   400  DKFGFIKSDYIVGHSSNSKGSSE.ETEAQYKRQEEVLRKFRHGEL
CEDCRWT                413  K.FKFVNPDYVVGASGRNLASSD..SQGLHKRQTEVLRRFHRNEI
ANCD1LOPHDEUT          430  D.LSFIKSSYITGHGAGSTGTSSKETEMQFRKQEEVLRKFRRHET
ANCD1DEUT              447  D.LSFISSSYITGHGAGSTGTSSKETEMQFRKQEEVLRKFRRHET
ANCD1DEUTSPECIES       447  D.LSFISSSYIVGHGSGSTGTSSKETEMQFRKQEEVLRKFRRHET
ANCD1VERT              460  E.LSYISSNFITGHGIGKNQPRNKQMEVEFRKQEEVLRKFRAHET
HSDCR                  470  E.LAYISSNFITGHGIGKNQPRNKQMEAEFRKQEEVLRKFRAHET
```

Motif IVa — Hel2 loop — Motif IVb (loop helix)

C

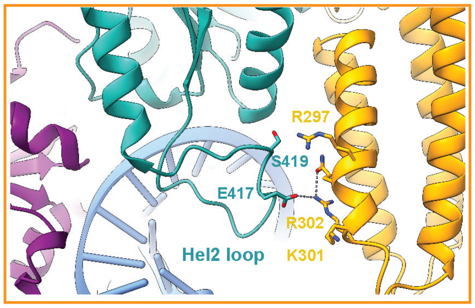


D

```
                                S415  T420
DMDCR2                 427  R...NNISPDFESVLERKWQKSAIQQFR
ANCD2ARTH              411  HNS.SPSSKEKSTEMSSKKQKEVLKKFR
ANCD1D2                411  HNS.SPSSKEKSTEMSSKKQKEVLKKFR
ANCD1D2SPECIES         409  HNS.SPSSKEKSTEMNSKKQKEVLKKFR
ANCD1BILATSPECIES      441  HGSGSTGTSSKETEMQFKKQEEVLRKFR
ANCD1NEMATODESPECIES   413  HSSNSKGSSE.ETEAQYKRQEEVLRKFR
CEDCRWT                425  ASGRNLASSD..SQGLHKRQTEVLRRFH
ANCD1ARTHLOPHDEUT      436  HGTGSTGTSSKETEMQFRKQEEVLRKFR
ANCD1LOPHDEUT          442  HGAGSTGTSSKETEMQFRKQEEVLRKFR
ANCD1DEUT              459  HGAGSTGTSSKETEMQFRKQEEVLRKFR
ANCD1DEUTSPECIES       459  HGSGSTGTSSKETEMQFRKQEEVLRKFR
AncD1VERT              472  HGIGKNQPRNKQMEVEFRKQEEVLRKFR
HSDCR                  482  HGIGKNQPRNKQMEAEFRKQEEVLRKFR
RIG-1                  664  RG.....KTNQNTGMTLPAQKCILDAFK.
DRH-1                  738  LN..KSTASSADISASKQKQMEKLKMFA
LGP2                   406  AG.....NSSQSTHMTQRDQQEVIQKFQ
                                T667  T671
```

Hel2 loop

**Fig. S15. Hel2 loop flexibility is important for helicase function.**
**(A)** Zoom-in view of the RecA-dsRNA interface comparing **state D.3** (Hel1, purple; Hel2, sea green) and **state D.4** (Hel1, pink; Hel2, blue), showing Hel2 motifs in proximity to dsRNA tracking strand. dsRNA is colored gray with orange and red for phosphate backbone; 3' and 5' directions indicated. Key protein residues and corresponding motifs are highlighted. Lines show connection between P3, H409 of IVa and Q427 of IVb in state D.3. These contacts are absent in state D.4. Flexible movement of Hel2 loop as it relaxes to ground state affects position of flanking motifs IVa and IVb.
**(B)** Multiple sequence alignment of ancient and modern Dicer helicase-DUF283 constructs illustrated with ESPRIPT(64), showing a portion of Hel2 containing the loop helix and the connected Hel2 loop. Motifs IVa (containing H409) and IVb (containing Q427 in the loop helix) flank Hel2 loop. H409 is conserved in all active helicases or replaced by an arginine in the modern dmDcr2. Inverted blue triangles indicate residues involved in ATPase and translocation activity. Red shading/white text indicates identity, no shading/red text indicates similarity, and black text indicates no conservation. Columns with black and red text have at least 70% conservation, represented by red text, while black text indicates the non-conserved or variant amino acids. Q427 is conserved in all displayed helicases. Extant proteins: CEDCRWT, *Caenorhabditis elegans DCR-1*; DMDCR2, *Drosophila melanogaster* Dicer-2; HSDCR, *Homo sapiens* Dicer. Ancestors (ANC) are from the Maximum likelihood gene tree unless designated as SPECIES. ANCD1D2SPECIES is AncD1D2 used in this paper, generated from species tree of life.
**(C)** Zoom-in view of the Hel2 loop interface with Hel2i in **state D**$^{post,ground}$. dsRNA is colored cornflower blue. Key protein residues are highlighted. Interactions between sidechains from Hel2 loop and Hel2i are depicted by dashed lines. S419 is in a cluster of conserved serines and threonines in active Dicer and RLR helicases. Hel2 loop flexibility likely creates different connections with Hel2i as translocation progresses.
**(D)** Multiple sequence alignment showing Hel2 loop from Dicer helicases described in (B) and the RIG-I-Like helicases, RIG-I, DRH-1 and LGP2. Red shading/white text indicates identity, no shading/red text indicates similarity, and black text indicates no conservation. Columns with black and red text have at least 70% conservation, represented by red text, while black text indicates the non-conserved or variant amino acids. Blue triangles indicate threonine residues essential for RIG-I translocation and signaling (16). Hel2 loop sequence shows low conservation but RIG-I T667 and T671 are either conserved or replaced with serine at the same position/adjacent positions (S414/415 and S419/T420 in AncD1D2SPECIES) in all functional Dicer and RLR helicases. Vertebrate Dicer helicases lack conservation at this site. Abbreviations as in (B).

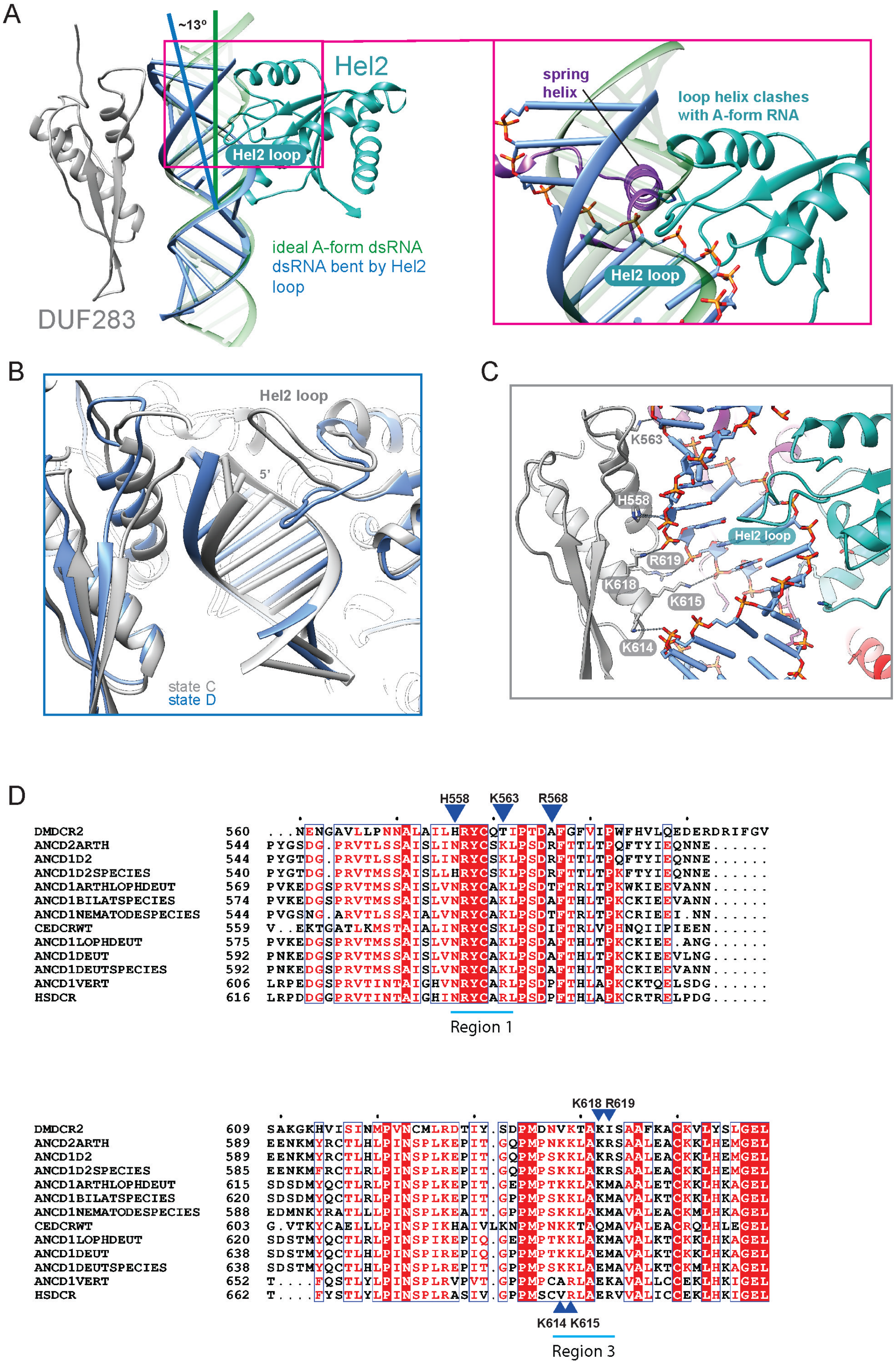
A
~13°
Hel2
Hel2 loop
ideal A-form dsRNA
dsRNA bent by Hel2
loop
DUF283
spring
helix
loop helix clashes
with A-form RNA
Hel2 loop
B
Hel2 loop
5'
state C
state D
C
K563
H558
Hel2 loop
R619
K618
K615
K614
D
H558
K563
R568
DMDCR2 560 ...NENGAVLLPNNALAILHRYCQTIPTDAFGFVIPWFHVLQEDERDRIFGV
ANCD2ARTH 544 PYGSDG.PRVTLSSAISLINRYCSKLPSDRFTTLTPQFTYIEQNNE......
ANCD1D2 544 PYGTDG.PRVTLSSAISLINRYCSKLPSDRFTTLTPQFTYIEQNNE......
ANCD1D2SPECIES 540 PYGTDG.PRVTMSSAISLLHRYCSKLPSDRFTTLTPKFTYIEQNNE......
ANCD1ARTHLOPHDEUT 569 PVKEDGSPRVTMSSAISLVNRYCAKLPSDTFTRLTPKWKIEEVANN......
ANCD1BILATSPECIES 574 PVKEDGSPRVTMSSAISLVNRYCAKLPSDAFTHLTPKCKIEEVANN......
ANCD1NEMATODESPECIES 544 PVGSNG.ARVTLSSAIALVNRYCAKLPSDTFTRLTPKCRIEEI.NN......
CEDCRWT 559 V..EKTGATLKMSTAIALINRYCSKLPSDIFTRLVPHNQIIPIEEN......
ANCD1LOPHDEUT 575 PVKEDGSPRVTMSSAISLVNRYCAKLPSDAFTHLTPKCKIEE.ANG......
ANCD1DEUT 592 PNKEDGSPRVTMSSAISLVNRYCAKLPSDAFTHLTPKCKIEEVLNG......
ANCD1DEUTSPECIES 592 PNKEDGSPRVTMSSAISLVNRYCAKLPSDAFTHLTPKCKIEEVANN......
ANCD1VERT 606 LRPEDGSPRVTINTAIGHVNRYCARLPSDPFTHLAPKCKTQELSDG......
HSDCR 616 LRPDDGGPRVTINTAIGHINRYCARLPSDPFTHLAPKCRTRELPDG......
Region 1
K618 R619
DMDCR2 609 SAKGKHVISINMPVNCMLRDTIY.SDPMDNVKTAKISAAFKACKVLYSLGEL
ANCD2ARTH 589 EENKMYRCTLHLPINSPLKEPIT.GQPMPNKKLAKRSAALEACKKLHEMGEL
ANCD1D2 589 EENKMYRCTLHLPINSPLKEPIT.GQPMPSKKLAKRSAALEACKKLHEMGEL
ANCD1D2SPECIES 585 EENKMFRCTLRLPINSPLREPIT.GQPMPSKKLAKRSAALEACKKLHEMGEL
ANCD1ARTHLOPHDEUT 615 SDSDMYQCTLRLPINSPLKEPIT.GEPMPTKKLAKMAAALETCKKLHKAGEL
ANCD1BILATSPECIES 620 SDSDMYQCTLRLPINSPLKEPIT.GPPMPSKKLAKMAVALKTCKKLHKAGEL
ANCD1NEMATODESPECIES 588 EDMNKYRATLLLPINSPLKEAIT.GPPMPSKKLAKMAVALEACKMLHKAGEL
CEDCRWT 603 G.VTKYCAELLLPINSPIKHAIVLKNPMPNKKTAQMAVALEACRQLHLEGEL
ANCD1LOPHDEUT 620 SDSTMYQCTLRLPINSPIKEPIQ.GEPMPTKKLAKMAVALKTCKKLHKAGEL
ANCD1DEUT 638 SDSTMYQCTLHLPINSPIREPIQ.GPPMPTKKLAEMAVALKTCKKLHKAGEL
ANCD1DEUTSPECIES 638 SDSTMYQCTLHLPINSPLREPIT.GPPMPSKKLAEMAVALKTCKMLHKAGEL
ANCD1VERT 652 T....FQSTLYLPINSPLRVPVT.GPPMPCARLAEKAVALLCCEKLHKIGEL
HSDCR 662 T....FYSTLYLPINSPLRASIV.GPPMSCVRLAERVVALICCEKLHKIGEL
K614 K615
Region 3

**Fig. S16. Hel2 loop distorts dsRNA structure and enables extensive contact with DUF283.**
**(A)** Front-view of colored model of AncD1D2 bound to 27-bp dsRNA in **state D**$^{post,ground}$ showing Hel2 (sea green) and DUF283 (gray). Hel2 loop is inserted into expanded dsRNA major groove (blue) compared with predicted structure of ideal A-form dsRNA (forest green). Helicase bound dsRNA is bent towards DUF283 (gray) during translocation in internal dsRNA segments. Inset: zoom-in view showing predicted clash between loop helix and A-form dsRNA during translocation.
**(B)** Zoom-in view of overlaid **state C**$^{post,closed}$ (gray) and **state D**$^{post,ground}$ (blue) models showing Hel2 loop transition from terminus capping position to major groove insertion position.
**(C)** Zoom-in view of extensive DUF283-dsRNA interface enabled by Hel2 loop (sea green) inserted into dsRNA major groove of **state D**$^{post,ground}$. Contacts between DUF283 (gray) and dsRNA backbone (blue with yellow and red for $PO_4$) are highlighted.
**(D)** Multiple sequence alignment of ancestral and modern Dicer Helicase-DUF283 constructs illustrated with ESPRIPT (64). Shown is the DUF283 segment depicting conservation of DUF283-dsRNA contact sites. Red shading/white text indicates identity, no shading/red text indicates similarity, and black text indicates no conservation. Columns with black and red text have at least 70% conservation, represented by red text, while black text indicates the non-conserved or variant amino acids. Blue triangles indicate residues involved in contact between DUF283 and dsRNA backbone in state E$^{internal‡}$. Regions 1 and 3 are canonical contact sites between the dsRBM fold and internal dsRNA segments (65). Conservation of contact residues is sporadic with loss of K618 and R619 correlating with decline in efficiency of dsRNA binding and translocation. K615 is conserved in all active helicases. K614 is replaced with V in dmDcr-2 but is otherwise conserved in all active ancestral helicases. Abbreviations as in S15B.

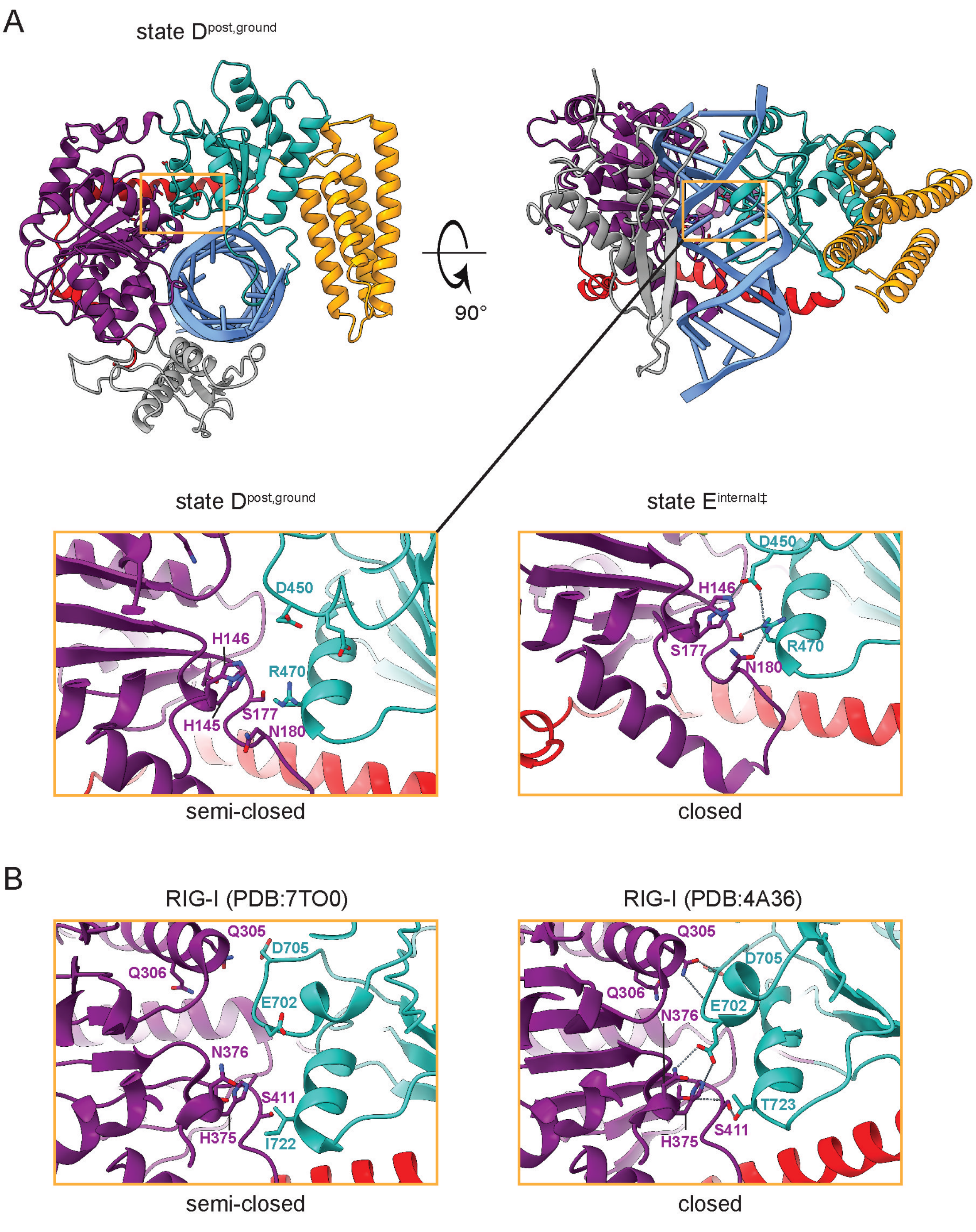


**Fig. S17. Structural comparison of Hel1-Hel2 interface between AncD1D2 and RIG-I. (A)** Top view (left) and front view (right) of colored atomic model of state $D^{post,ground}$. Inset shows Hel1-Hel2 interface in state $D^{post,ground}$ (left) and state $E^{internal‡}$ (right). **(B)** RIG-I Hel1-Hel2 interface showing semi-closed (PDB:7TO0) and closed (PDB:4A36) conformations. Equivalent residues between AncD1D2 and RIG-I are indicated, H145:H375, H146:N376, S177:S411, D450:E702.

**Table S1.** Parameters for ATP hydrolysis, dsRNA affinity and translocation

| **Condition (RNA)** | **$k_{burst}$ (μM/min)** | **$k_{obs(TLC)}$ ($min^{-1}$)** | **$k_{cat}$ ($min^{-1}$)** | **$K_M$ (μM)** | **$k_{cat}/K_M$ ($μM^{-1}$ $min^{-1}$)** | **$K_D$ (nM)** | **$k_{obs(STREP)}$ ($min^{-1}$)** | **Amplitude (% strep displaced)** |
|---|---|---|---|---|---|---|---|---|
| AncD1D2 (none) | -- | 0.07 ± 0.03 | | | | | | |
| AncD1D2 (BLT) | 19.7 ± 3.5 | 0.18 ± 0.04 | 358 ± 51 | 293 ± 128 | 1.22 | 76 | 0.20 ± 0.05 | 57 |
| AncD1D2 (3'ovr) | 17.6 ± 1.3 | 0.24 ± 0.03 | 203 ± 30 | 106 ± 62 | 1.91 | | 0.18 ± 0.06 | 49 |
| $AncD1_{BILAT}$ (none) | -- | 0.08 ± 0.04 | | | | | | |
| $AncD1_{BILAT}$ (BLT) | 13.3 ± 3.4 | 0.17 ± 0.02 | 172 ± 20 | 284 ± 104 | 0.61 | 140 | 0.03 ± 0.03 | 30 |
| $AncD1_{BILAT}$ (3'ovr) | 11.8 ± 3.6 | 0.12 ± 0.01 | 73 ± 10 | 126 ± 72 | 0.57 | | 0.03 ± 0.03 | 25 |
| $AncD1_{DEUT}$ (none) | -- | 0.07 ± 0.02 | | | | | | |
| $AncD1_{DEUT}$ (BLT) | 10.0 ± 1.0 | 0.09 ± 0.02 | 76 ± 9 | 114 ± 52 | 0.66 | 965 | u.d. | |
| $AncD1_{DEUT}$ (3'ovr ) | 5.8 ± 1.2 | 0.07 ± 0.02 | 129 ± 23 | 700 ± 296 | 0.18 | | u.d. | |
| $AncD1_{NEM}$ (none) | -- | 0.04 ± 0.01 | | | | | | |
| $AncD1_{NEM}$ (BLT) | 1.5 ± 0.9 | 0.04 ± 0.02 | | | | | u.d. | |
| $AncD1_{NEM}$ (3'ovr) | 1.2 ± 1.0 | 0.04 ± 0.02 | | | | | u.d. | |

u.d. – undetectable

**Table S2.** Cryo-EM data collection, refinement and validation statistics

| Structure | State A | State B | State C | State D | State E |
|---|---|---|---|---|---|
| EMDB accession ID | EMD-48678 | EMD-48691 | EMD-48697 | EMD-48708 | EMD-48710 |
| PDB accession ID | 9MW6 | 9MW7 | 9MW8 | 9MX3 | 9MX5 |
| **Data collection and processing** | | | | | |
| Microscope | Titan Krios G3 | | | | |
| Voltage (kV) | 300 | | | | |
| Detector | Gatan K3 | | | | |
| Nominal magnification | 105,000X | | | | |
| Total number of frames | 50 | | | | |
| Total electron exposure ($e^-$/Å$^2$) | 40 | | | | |
| Defocus range (μM) | -0.8 to -2.4 | | | | |
| Number of micrographs | 11,003 | 11,834 | 15,138 | 15,138 | 7,335 |
| Pixel size (Å) | 0.436 (super-resolution) | 0.872 | 0.436 (super-resolution) | 0.436 (super-resolution) | 0.436 (super-resolution) |
| Symmetry imposed | C1 | C1 | C1 | C1 | C1 |
| Initial number of particles | 12,548,917 | 23,429,138 | 21,532,216 | 21,532,216 | 13,806,533 |
| Final number of particles | 303,540 | 380,191 | 315,941 | 510,474 | 171,095 |
| Map resolution (masked, corrected) (Å) | 3.4 | 3.4 | 3.3 | 3.0 | 3.1 |
| FSC threshold | 0.143 | 0.143 | 0.143 | 0.143 | 0.143 |
| Map resolution range (Å) | 2.0 - 11.9 | 2.0 - 41.0 | 1.9 - 8.7 | 1.9 - 31.8 | 1.9 - 7.0 |
| **Refinement** | | | | | |

| | | | | | |
|---|---|---|---|---|---|
| Initial model used | AlphaFold 2 | AlphaFold 2 | AlphaFold 2 | AlphaFold 2 | AlphaFold 2 |
| Model resolution (FSC = 0.5) (Å) | 4.2 | 4.4 | 4.2 | 3.4 | 3.5 |
| Model composition | | | | | |
| Non-hydrogen atoms | 5,473 | 5,504 | 6,419 | 5,929 | 6,463 |
| Protein residues | 535 | 535 | 647 | 641 | 652 |
| RNA residues | 54 | 54 | 54 | 33 | 54 |
| Ligands<br>ADP/AF3/Mg/ | 0/0/0 | 1/1/0 | 1/0/1 | 1/0/0 | 1/1/1 |
| R.m.s deviations | | | | | |
| Bond lengths (Å) | 0.002 | 0.002 | 0.002 | 0.003 | 0.003 |
| Bond angles (°) | 0.566 | 0.619 | 0.634 | 0.634 | 0.555 |
| Validation | | | | | |
| Molprobity score | 1.92 | 2.25 | 1.96 | 1.81 | 1.91 |
| Clashscore | 10.04 | 16.09 | 13.58 | 6.90 | 10.04 |
| Poor/Outlier rotamers (%) | 0.41 | 1.01 | 1.33 | 1.68 | 1.32 |
| Ramachandran plot | | | | | |
| Favored (%) | 94.16 | 90.58 | 96.59 | 96.24 | 95.69 |
| Allowed (%) | 5.84 | 9.23 | 3.26 | 3.76 | 4.15 |
| Outliers (%) | 0.00 | 0.19 | 0.16 | 0.00 | 0.15 |

**Table S3.** Extant and ancestrally reconstructed Dicer helicase sequences

| **Name** | **Sequence** |
|---|---|
| DMDCR2<br><br>*Drosophila melanogaster* Dicer-2 | MEDVEIKPRGYQLRLVDHLTKSNGIVYLPTGSGKTFVAIL<br>VLKRFSQDFDKPIESGGKRALFMCNTVELARQQAMAVRRC<br>TNFKVGFYVGEQGVDDWTRGMWSDEIKKNQVLVGTAQVFL<br>DMVTQTYVALSSLSVVIIDECHHGTGHHPFREFMRLFTIA<br>NQTKLPRVVGLTGVLIKGNEITNVATKLKELEITYRGNII<br>TVSDTKEMENVMLYATKPTEVMVSFPHQEQVLTVTRLISA<br>EIEKFYVSLDLMNIGVQPIRRSKSLQCLRDPSKKSFVKQL<br>FNDFLYQMKEYGIYAASIAIISLIVEFDIKRRQAETLSVK<br>LMHRTALTLCEKIRHLLVQKLQDMTYDDDDDNVNTEEVIM<br>NFSTPKVQRFLMSLKVSFADKDPKDICCLVFVERRYTCKC<br>IYGLLLNYIQSTPELRNVLTPQFMVGRNNISPDFESVLER<br>KWQKSAIQQFRDGNANLMICSSVLEEGIDVQACNHVFILD<br>PVKTFNMYVQSKGRARTTEAKFVLFTADKEREKTIQQIYQ<br>YRKAHNDIAEYLKDRVLEKTEPELYEIKGHFQDDIDPFTN<br>ENGAVLLPNNALAILHRYCQTIPTDAFGFVIPWFHVLQED<br>ERDRIFGVSAKGKHVISINMPVNCMLRDTIYSDPMDNVKT<br>AKISAAFKACKVLYSLGELNERFVPKTLKERVA |
| AncD2$_{ARTH}$<br><br>Ancestor of Arthropod Dicer-2 from gene tree | MDETDEDEFTPRPYQVELLERAMKKNTIVCLGTGSGKTFI<br>AVMLIKELAHEIRGPFSEGGKRTIFLVNTVPLVNQQAKVI<br>RRHTSLKVGEYTGDMNVDSWNKEKWNQEFEKHQVLVMTAQ<br>IFLDILNHGFISLNQINLLIFDECHHAVKNHPMRQIMRHY<br>KNLEQNDRPRILGLTASVINSKCKPNQVEKKIKELETTMN<br>SKVVTASDLEEVAVQKYSTKPKEIIVSYDNDRQSDTSEVI<br>ENIINQALEQLSNIEETSNLNDTNSLKQIKKVLRDIKNIL<br>DELGPWCAHRVIKSRIEQLEKRESETAEELRTIRELLQSIF<br>EQIINVLKNLEKLQKIKNNSVEYVSPKVRKLLEILKQYYSNNN<br>NSKEELCGIIFVERRYTAKVLYHLLKELSKKHDDDFS<br>FIKPDFIVGHNSSPSSKEKSTEMSSKKQKEVLKKFRKGEC<br>NLLVATSVLEEGIDIPKCNLVVRFDLPKNFRSYVQ<br>SKGRARAKNSHYIIMVEEDEKNKFLEDLNQYQEIEKILL<br>RLCHNRNAPTSEEDFDNFEVDDLLPPYMPYGSDGPR<br>VTLSSAISLINRYCSKLPSDRFTTLTPQFTYIEQNNEEEN<br>KMYRCTLHLPINSPLKEPITGQPMPNKKLAKRSAALEACK<br>KLHEMGELDDHLLPVSISRKNAELK |
| AncD1D2<br><br>Pre-gene duplication ancestor of metazoan Dicer from gene tree | MDETDEDEFTPRPYQVELLERAMKKNTIVCLGTGSGKTFI<br>AVMLIKELAHEIRGPFSEGGKRTFFLVNTVPLVNQQAKVI<br>RKHTSLKVGEYVGDMGVDSWNKEKWNQEFEKHQVLVMTAQ<br>IFLDILNHGFISLNQINLLIFDECHHAVKNHPYRQIMRHY<br>KNLEQNDRPRILGLTASVINSKCKPNQVEKKIKELEATLN<br>SRVVTASDLEEVAVQKYATKPKEIIVSYNNDRKSDTSEVI<br>ENIINQALEQLSNIEETSNLNDTNSLKQIKKVLRDIKNIL<br>DELGPWCAHRVIKSRIRQLEKRESETAEELRTIRELLQSI<br>FEQIINVLKNLEKLQKIKNKSVEYVSPKVRKLLEILKQYF<br>SNNNNSNEELCGIIFVERRYTAYVLYKLLKELSKKHDDDF<br>SFIKCDFIVGHNSSPSSKEKSTEMSSKKQKEVLKKFRKGE<br>LNLLVATSVVEEGIDIPKCNLVVRFDLPKNFRSYVQSKGR<br>ARAKNSHYIIMVEEDEKNKFQEDLNQYQEIEKILLRLCHN<br>RNAPTSEEDFDNFEVDDLLPPYMPYGTDGPRVTLSSAISL<br>INRYCSKLPSDRFTTLTPQFTYIEQNNEEENKMYRCTLHL<br>PINSPLKEPITGQPMPSKKLAKRSAALEACKKLHEMGELD<br>DHLLPVSISRKNAELK |

| | |
|---|---|
| AncD1D2$_{SPECIES}$<br><br>Pre-gene duplication ancestor of metazoan Dicer from species tree | MDETDEDEFTPRPYQVELLERAMKKNTIVCLGTGSGKTFI<br>AVMLIKELAHEIRGPFNEGGKRTFFLVNTVPLVNQQAKVI<br>RKHTSLKVGEYVGDMGVDSWNKEKWNQEFEKHQVLVMTAQ<br>IFLDILNHGFISLSQVNLLIFDECHHAVKNHPYRQIMRHY<br>KNLEQNDRPRILGLTASVINSKCKPNQVEKKIKELEATLN<br>SRVVTASDLEEVAVQKYATKPKEIIVSYNSDRKSDTSEVI<br>ENIINQALEQLSNIEETSNLNDTNSLKQIKKVLRDIKNIL<br>DELGPWCAHRVIKSRIRQLEKRESETAEELRTIRELLQSI<br>FEQIINVLKNLEKLQKNNSVEFVSPKVKKLLEILKQYFSN<br>NNNSSKELCGIIFVERRYTAYVLYKLLNELSAKRDDDFSF<br>IKCDFVVGHNSSPSSKEKSTEMNSKKQKEVLKKFRKGECN<br>LLVATSVVEEGIDIPKCNLVVRFDLPKNFRSYVQSKGRAR<br>AKNSKYIIMVEEDEKNKFQEDLNQYQEIEKILLRLCHNRD<br>APSEEDFDSFEDELLPPYMPYGTDGPRVTMSSAISLLHRY<br>CSKLPSDRFTTLTPKFTYIEQNNEEENKMFRCTLRLPINS<br>PLREPITGQPMPSKKLAKRSAALEACKKLHEMGELDDHLL<br>PVKISRKNAELK |
| AncD1$_{ARTHLOPHDEUT}$<br><br>Common ancestor of arthropod, lophotrochozoan and deuterostome Dicer-1 from gene tree | MPSGMSKPLENIHTNTFTPRPYQVELLDAAKQRNTIVCLG<br>TGTGKTFIAVMLIKELAHQIRRPFNDGGKRTFFLVNTVPL<br>VSQQAKVIRHHTDLSVGEYVGDMDVDSWNKEKWKQEFEKH<br>QVLVMTAQIFLDILQHGFLSLSKVNLLIFDECHHAVKNHPYRQI<br>MKMFDNCPQNNRPRILGLTASLLNSKCKPNQLEKKIRELE<br>KTLRSTVETASDLVSVSRYGTKPKEIVVEYNSSDSDEEDD<br>TELSNEIQNILNDALAFLNDNIEEEENISDPCVQPKKVLN<br>ECLYILNELGPWCADRVAQMFIKEIEKLETKHVSSEIHRL<br>FLQYTHTQLRMIRICENAFKESENVEKLLKFVSPKVRRLL<br>EILKEYKPSSESDENDNSQQQESNSEDESDSLCGIIFVER<br>RYTAYVLNKLLKELSKRDPDFSFIKSDYIVGHGTGSTGTS<br>SKETEMQFRKQEEVLRKFRKHECNLLVATSVVEEGVDVPK<br>CNLVVRFDLPKNYRSYVQSKGRARAPDSHYIMLVEEDEKE<br>KFQEDLKNYHEIEKILLRKCHDREAPTEEEIDASIADNLI<br>PPYMPVKEDGSPRVTMSSAISLVNRYCAKLPSDTFTRLTP<br>KWKIEEVANNSDSDMYQCTLRLPINSPLKEPITGEPMPTK<br>KLAKMAAALETCKKLHKAGELDDHLLPVGKETIKDE |
| AncD1$_{BILATSPECIES}$<br><br>Common ancestor of bilaterian Dicer-1 from species tree | MPSGMSKPLQNENIHTNTFTPRPYQVELLDAAKQKNTIVC<br>LGTGTGKTFIAVMLIKELAHQIRRPLNDGGKRTFFLVNSV<br>PLVSQQAKVIRHHTDLNVGEYVGADMDVDSWNKEKWNQEF<br>EKHQVLVMTAQIFLDILQHGFLSLSKVNLLIFDECHHAVK<br>NHPYRQIMKMFDNCPQNNRPRILGLTASILNSKCKPNQLE<br>KKIRELEKTLRSTAETASDLVSVSRYGTKPKEVIVECSDS<br>YQEDDTELSNEIDNILNDALAFLNDCNIALEEERDPCAIP<br>KKVLNECLYILNQLGPWCANRVAQMFIKELEKLETKHESS<br>EIHRLFLQYTQTQLRMIRKICENAFKESENVEKLLKFVSP<br>KVRRLLEILKEYKPSSEEDENDNSQQQESSRDNSEDEDDS<br>LCGIVFVERRYTAYVLNKLLKELSKRDPDFSFIKSDYIVG<br>HGSGSTGTSSKETEMQFKKQEEVLRKFRRHECNLLVATSV<br>VEEGVDVPKCNLVVRFDLPKNYRSYVQSKGRARAKNSHYI<br>MLVEEDEMEKFQEDLKNYQEIEKILLRKCHDREEPDEEEI<br>DASIADNLIPPYMPVKEDGSPRVTMSSAISLVNRYCAKLP<br>SDAFTHLTPKCKIEEVANNSDSDMYQCTLRLPINSPLKEP<br>ITGPPMPSKKLAKMAVALKTCKKLHKAGELDDHLLPVGKE<br>TIKYE |
| AncD1$_{NEMATODESPECIES}$ | MMSKSSDVENDFFTPRDYQVELLDKAKKRNTIVPLGTGSG<br>KTFIAVMLIKELAAQIRRPLSEGGKRTFFVVDKVPLVEQQ<br>AEHIRHHTDLNVGEFHGDLNVDSWNSEKWNTFFEEHQVLV<br>MTAQIFLDLLNHGFLNLDNINLLIFDECHHALKNHPYRQI |

| | |
|---|---|
| Common ancestor of nematode Dicer-1 from species tree | MKRYSKLPNENRPRILGLTASLINDKVKPNQLEKKIRKLE<br>RTLHSKVETASDLVSVSKYGAKPKEVVIVCRDFDTSNECL<br>DNILQLIEELEKLCKTSDLDVNCMKHIKEALNKILSILNQ<br>LGPWCAWKVCQKRERQLKKLEKRITLSETQHLFLQMGQTT<br>LRTVRKLLEPKVKNVKSVEELKPFVSNKVRRLLEILETYN<br>KRSLENDNNESLCGIIFVEQRYVAYVLNILLKELSKWDPD<br>KFGFIKSDYIVGHSSNSKGSSEETEAQYKRQEEVLRKFRH<br>GELNLLVATSVLEEGIDVRQCNLVIRFDLPTNFRSYVQSK<br>GRARKRDAHYIILVEEKDSENFQEDLKNFVEIEKILLRRC<br>HNSSENVELRYENVDEDLIPPYVPVGSNGARVTLSSAIAL<br>VNRYCAKLPSDTFTRLTPKCRIEEINNEDMNKYRATLLLP<br>INSPLKEAITGPPMPSKKLAKMAVALEACKMLHKAGELND<br>HLLPVGKETIKKV |
| CEDCRWT<br><br>*Caenorhabditis elegans* Dicer-1 | MVRVRADLQCFNPRDYQVELLDKATKKNTIVQLGTGSGKT<br>FIAVLLLKEYGVQLFAPLDQGGKRAFFVVEKVNLVEQQAI<br>HIEVHTSFKVGQVHGQTSSGLWDSKEQCDQFMKRHHVVVI<br>TAQCLLDLIRHAYLKIEDMCVLIFDECHHALGSQHPYRSI<br>MVDYKLLKKDKPVPRVLGLTASLIKAKVAPEKLMEQLKKL<br>ESAMDSVIETASDLVSLSKYGAKPYEVVIICKDFEIGCLG<br>IPNFDTVIEIFDETVAFVNTTTEFHPDLDLDPRRPIKDSL<br>KTTRAVFRQLGPWAAWRTAQVWEKELGKIIKSQVLPDKTL<br>RFLNMAKTSMITIKRLLEPEMKKIKSIEALRPYVPQRVIRL<br>FEILETFNPEFQKERMKLEKAEHLSAIIFVDQRYIAYSLL<br>LMMRHIKSWEPKFKFVNPDYVVGASGRNLASSDSQGLHKR<br>QTEVLRRFHRNEINCLIATSVLEEGVDVKQCNLVIKFDRP<br>LDMRSYVQSKGRARRAGSRYVITVEEKDTAACDSDLKDFQ<br>QIEKILLSRHRTVNNPIEDDSDRFEEFDVDSQMEPYVVEK<br>TGATLKMSTAIALINRYCSKLPSDIFTRLVPHNQIIPIEE<br>NGVTKYCAELLLPINSPIKHAIVLKNPMPNKKTAQMAVAL<br>EACRQLHLEGELDDNLLPKGRESI |
| AncD1$_{LOPHDEUT}$<br><br>Common ancestor of lophotrochozoan and deuterostome Dicer-1 from gene tree | MPSGSSKPLQNPRQENIHTNTFTPRPYQVELLDAALQQNT<br>IVCLGTGTGKTFIAVMLIKELAHQIRRPLNDGGKRTFFLV<br>NSVPLVSQQAKVIRHHTDLSVGEYVGAMDVDSWNKEKWNQ<br>EFEKHQVLVMTAQIFLDILQHGFLSLSKVNLLIFDECHHA<br>VKNHPYRQIMKMFDNCPKNNRPRILGLTASILNSKCKPNE<br>LEKKIRELEKTLRSTAETATDLVSVDRYGTKPKEVVIECD<br>SYEDDTGLVNEINNILNDALAFLNDCNIALEEEERDPCVI<br>PKQVLNECLVILNELGPWCADRVAQMFIKELEKLEKHESS<br>EIHRLFLQYTQTQLRMIHKICENAFKESENVEKLLKFVTP<br>KVRRLLEILKEYKPSSEEDENDSSQQQESSRQDNSEDEDD<br>SLCGIVFVERRYTAFVLNKLLKELSKRDPDLSFIKSSYIT<br>GHGAGSTGTSSKETEMQFRKQEEVLRKFRRHETNLLVATS<br>VVEEGVDVPKCNLVVRFDLPKNYRSYVQSKGRARAPDSHY<br>IMLVEEDELEAFQEDLKNYKEIEKILLRKCHEREEPTEEE<br>IDASIADNLLPPYMPVKEDGSPRVTMSSAISLVNRYCAKL<br>PSDAFTHLTPKCKIEEANGSDSTMYQCTLRLPINSPIKEPI<br>QGEPMPTKKLAKMAVALKTCKKLHKAGELDDHLLPVGKET<br>IKYE |
| AncD1$_{DEUT}$<br><br>Common ancestor of deuterostome Dicer-1 from gene tree | MLPSGSSKPLQNENIHTNTFTPRPYQVELLDAALEQNTIV<br>CLGTGTGKTFIAVMLIKELSHQIRRPLNDGGKRTIFLVNS<br>VPLVSQQAKVIRTHTDLSVGEYVGAMDVDSWNKEKWNQEF<br>EKHQVLVMTAQIFLDILQHGFLSLSRVNLLIFDECHHAVK<br>NHPYREIMKMFDNCPKNNRPRILGLTASILNGKCKPNELE<br>KKIRELEKTLRSTAETATDLVSVDRYGTKPKEVVIECESY<br>EDDTGLSSEINNILNDALAFLNDCNISLEEEERDPCLIPK<br>QVLNECLVILNVLGPWCANRVAQMLIKELEKLEKHESSEI |

| | |
|---|---|
| | HRLFLQYTQTQLRMIHKICENAFKESENVDLKFVTPKVRR<br>LLEILKEYKPSEEDENDSSQQQESRHNNRNDNYVSEDEDD<br>DSESEEEEEKQDTNSTTSLCGIVFVERRYTAVVLNKLLKE<br>LSKRDPDLSFISSSYITGHGAGSTGTSSKETEMQFRKQEE<br>VLRKFRRHETNLLVATSVVEEGVDVPKCNLVVRFDLPKNY<br>RSYVQSKGRARAPNSHYIMLVEEDELEAFKEDLKNYKEIE<br>KILLRKCHEREEPQEEEIDAHIADNLLPPYMPNKEDGSPR<br>VTMSSAISLVNRYCAKLPSDAFTHLTPKCKIEEVLNGSDS<br>TMYQCTLHLPINSPIREPIQGPPMPTKKLAEMAVALKTCK<br>KLHKAGELDDHLLPVGKETIKYE |
| AncD1$_{DEUTSPECIES}$<br><br>Common ancestor of deuterostome Dicer-1 from species tree | MLPSGSSKPLQNENIHTNTFTPRPYQVELLDAALEKNTIV<br>CLGTGTGKTFIAVMLIKELSHQIRRPLNDGGKRTIFLVNS<br>VPLVSQQANVIRTHTDLNVGEYVGAMDVDSWNKEKWNQEF<br>EKHQVLVMTAQIFLDILQHGFLSLSRVNLLIFDECHHAIK<br>NHPYRQIMKIFDNCPQNNRPRILGLTASILNGKCKPNQLE<br>KKIRELEKTLRSTAETATDLVSVSRYGTKPKEVVIECESY<br>EDDTGLSNEINNILNDALAFLNDCNISLEEEERDPCLIPK<br>QVLNECLAILNVLGPWCANRVAQMLIKELEKLEKHESSEI<br>HRLFLQYTQTQLRMVRKLCENAFKESENVDLKFVTPKVRR<br>LLEILKEYKPSEEDENDNSQQTESRHNNRNDNYVSEDEDD<br>DSESEEEEEKQDTNSTTSLCGIVFVERRYTAVVLNKLLKE<br>LSKRDPDLSFISSSYIVGHGSGSTGTSSKETEMQFRKQEE<br>VLRKFRRHETNLLVATSVVEEGVDVPKCNLVVRFDLPKNY<br>RSYVQSKGRARAPNSHYIMLVQEDEMEAFKEDLKNYQEIE<br>KILLRKCHDREEPQEEEIDAHIADNLLPPYMPNKEDGSPR<br>VTMSSAISLVNRYCAKLPSDAFTHLTPKCKIEEVANNSDS<br>TMYQCTLHLPINSPLREPITGPPMPSKKLAEMAVALKTCK<br>MLHKAGELDDHLLPVGKETIKYE |
| AncD1$_{VERT}$<br><br>Common ancestor of vertebrate Dicer-1 from gene and species trees | MAGLQLMTPASSPMGPFFGLPWQQEAIHDNIFTPRKYQVE<br>LLEAALEHNTIVCLNTGSGKTFIAVLLTKELSHQIRGQFN<br>KNGKRTVFLVNSAPSVAQQAAAVRTHSDLKVGEYSSLEDV<br>ESWTKEKWNQEFTEHQVLVMTCHIFLHILKNGFLSLSKIN<br>LLVFDECHLAIKDHPYREIMKICENCPSCPRILGLTASIL<br>NGKCDPSELEEKIQKLEKILRSNAETATDLVVLDRYSSQP<br>REVVLDCGPYVDKSGLYEKLLNELDEALNFLNDCNISVHS<br>EDRDPTLIPKQVLSDCRAVLTVLGPWCADKVAGMMVRELQ<br>KYIKHEQEELNRKFLLFTDTLLRKIHALCEEHFSPASLDL<br>KFVTPKVIKLLEILHEYKPFERQQFESVEWYNNRNQDNYV<br>SWSDSEDEDEDEEIEEKEKPDTNFPSPFTNILCGIIFVER<br>RYTAVVLNRLIKEAGKQDPELSYISSNFITGHGIGKNQPR<br>NKQMEVEFRKQEEVLRKFRAHETNLLIATSVVEEGVDIPK<br>CNLVVRFDLPTEYRSYVQSKGRARAPISNYIMLADSDKIE<br>AFKEDLKTYKAIEKILRNKCSKSADSSELEIEPVADDDDI<br>LPPYVLRPEDGSPRVTINTAIGHVNRYCARLPSDPFTHLA<br>PKCKTQELSDGTFQSTLYLPINSPLRVPVTGPPMPCARLA<br>EKAVALLCCEKLHKIGELDDHLMPVGKETVKYE |
| HSDCR<br><br>*Homo sapiens* Dicer-1 | MKSPALQPLSMAGLQLMTPASSPMGPFFGLPWQQEAIHDN<br>IYTPRKYQVELLEAALDHNTIVCLNTGSGKTFIAVLLTKE<br>LSYQIRGDFSRNGKRTVFLVNSANQVAQQVSAVRTHSDLK<br>VGEYSNLEVNASWTKERWNQEFTKHQVLIMTCYVALNVLK<br>NGYLSLSDINLLVFDECHLAILDHPYREIMKLCENCPSCP<br>RILGLTASILNGKCDPEELEEKIQKLEKILKSNAETATDL<br>VVLDRYTSQPCEIVVDCGPFTDRSGLYERLLMELEEALNF<br>INDCNISVHSKERDSTLISKQILSDCRAVLVVLGPWCADK<br>VAGMMVRELQKYIKHEQEELHRKFLLFTDTFLRKIHALCE<br>EHFSPASLDLKFVTPKVIKLLEILRKYKPYERQQFESVEW |

| | |
|---|---|
| | YNNRNQDNYVSWSDSEDDDEDEEIEEKEKPETNFPSPFTN<br>ILCGIIFVERRYTAVVLNRLIKEAGKQDPELAYISSNFIT<br>GHGIGKNQPRNKQMEAEFRKQEEVLRKFRAHETNLLIATS<br>IVEEGVDIPKCNLVVRFDLPTEYRSYVQSKGRARAPISNY<br>IMLADTDKIKSFEEDLKTYKAIEKILRNKCSKSVDTGETD<br>IDPVMDDDDVFPPYVLRPDDGGPRVTINTAIGHINRYCAR<br>LPSDPFTHLAPKCRTRELPDGTFYSTLYLPINSPLRASIV<br>GPPMSCVRLAERVVALICCEKLHKIGELDDHLMPVGKETV<br>KYE |

**Movie S1. Closure of Dicer helicase RecA domains is caused by ATP binding.** Transition from State A to State B is depicted. Labels as in Fig. 5.

**Movie S2. ATP hydrolysis triggers helicase movement in the closed state.** Transition from State B to State C is depicted. Interactions between V70 and H406 sidechains and phosphate backbone are depicted with dashed lines. Labels as in Fig. 5.

**Movie S3. Helicase transitions from post hydrolysis closed state to post hydrolysis semi-closed state.** Transition from State C to State D is depicted. Labels as in Fig. 5.

**Movie S4. Flexibility of Hel2 loop as revealed by from state D subclassification.** Movement of Hel2 loop affects position of flanking motifs: H409 (IVa) and Q427 (IVb).

**Movie S5. Model of Dicer helicase function depicting complete coupling of ATPase cycle to translocation on dsRNA.**

**SI References**


1. A. M. Aderounmu, P. J. Aruscavage, B. Kolaczkowski, B. L. Bass, Ancestral protein reconstruction reveals evolutionary events governing variation in Dicer helicase function. *Elife* **12**, e85120 (2023).

2. F. Lemoine, *et al.*, Renewing Felsenstein's Phylogenetic Bootstrap in the Era of Big Data. *Nature* **556**, 452–456 (2018).

3. A. M. Kozlov, D. Darriba, T. Flouri, B. Morel, A. Stamatakis, RAxML-NG: a fast, scalable and user-friendly tool for maximum likelihood phylogenetic inference. *Bioinformatics* **35**, 4453–4455 (2019).

4. N. K. Sinha, B. L. Bass, Overexpression and purification of Dicer and accessory proteins for biochemical and structural studies. *Methods* **126**, 54–65 (2017).

5. C. D. Consalvo, *et al.*, Caenorhabditis elegans Dicer acts with the RIG-I-like helicase DRH-1 and RDE-4 to cleave dsRNA. *eLife* **13**, RP93979 (2024).

6. A. Punjani, J. L. Rubinstein, D. J. Fleet, M. A. Brubaker, cryoSPARC: algorithms for rapid unsupervised cryo-EM structure determination. *Nat Methods* **14**, 290–296 (2017).

7. T. Bepler, *et al.*, Positive-unlabeled convolutional neural networks for particle picking in cryo-electron micrographs. *Nat. Methods* **16**, 1153–1160 (2019).

8. J. Jumper, *et al.*, Highly accurate protein structure prediction with AlphaFold. *Nature* **596**, 583–589 (2021).

9. P. Emsley, K. Cowtan, Coot: model-building tools for molecular graphics. *Acta Crystallogr. Sect. D: Biol. Crystallogr.* **60**, 2126–2132 (2004).

10. E. F. Pettersen, *et al.*, UCSF Chimera—A visualization system for exploratory research and analysis. *J. Comput. Chem.* **25**, 1605–1612 (2004).

11. T. D. Goddard, *et al.*, UCSF ChimeraX: Meeting modern challenges in visualization and analysis. *Protein Sci.* **27**, 14–25 (2018).

12. P. D. Adams, *et al.*, PHENIX: a comprehensive Python-based system for macromolecular structure solution. *Acta Crystallogr. Sect. D: Biol. Crystallogr.* **66**, 213–221 (2010).

13. R. Sanchez-Garcia, *et al.*, DeepEMhancer: a deep learning solution for cryo-EM volume post-processing. *Commun Biology* **4**, 874 (2021).

14. P. V. Afonine, *et al.*, Real-space refinement in PHENIX for cryo-EM and crystallography. *Acta Crystallogr. Sect. D* **74**, 531–544 (2018).

15. C. J. Williams, *et al.*, MolProbity: More and better reference data for improved all-atom structure validation. *Protein Sci.* **27**, 293–315 (2018).

16. S. C. Devarkar, B. Schweibenz, C. Wang, J. Marcotrigiano, S. S. Patel, RIG-I Uses an ATPase-Powered Translocation-Throttling Mechanism for Kinetic Proofreading of RNAs and Oligomerization. *Mol Cell* **72**, 355-368.e4 (2018).
